# Single-molecule visualization of stalled replication-fork rescue by the *Escherichia coli* Rep helicase

**DOI:** 10.1101/2022.12.04.519054

**Authors:** Kelsey S. Whinn, Zhi-Qiang Xu, Slobodan Jergic, Nischal Sharma, Lisanne M. Spenkelink, Nicholas E. Dixon, Antoine M. van Oijen, Harshad Ghodke

**Affiliations:** Molecular Horizons and School of Chemistry and Molecular Bioscience, University of Wollongong, Wollongong, New South Wales 2522, Australia; Illawarra Health & Medical Research Institute, Wollongong, New South Wales 2522, Australia

## Abstract

Genome duplication occurs while the template DNA is bound by numerous DNA-binding proteins. Each of these proteins act as potential roadblocks to the replication fork and can have deleterious effects on cells. In *Escherichia coli*, these roadblocks are displaced by the accessory helicase Rep, a DNA translocase and helicase that interacts with the replisome. The mechanistic details underlying the coordination with replication and roadblock removal by Rep remain poorly understood. Through real-time fluorescence imaging of the DNA produced by individual *E. coli* replisomes and the simultaneous visualization of fluorescently-labeled Rep, we show that Rep continually surveils elongating replisomes. We found that this association of Rep with the replisome is stochastic and occurs independently of whether the fork is stalled or not. Further, we visualize the efficient rescue of stalled replication forks by directly imaging individual Rep molecules as they remove a model protein roadblock, dCas9, from the template DNA. Using roadblocks of varying DNA-binding stabilities, we conclude that replication restart is the rate-limiting step of stalled replication rescue.

## INTRODUCTION

Cell proliferation requires high-fidelity duplication of the genome. In all cells, this process is achieved by a group of proteins, collectively known as the replisome. In *Escherichia coli*, more than 12 proteins coordinate the unwinding and duplication of DNA at rates of up to 1000 base pairs (bp) per second (1–5) and at processivities of 100s of kilobase pairs (kbp) (6–8). DNA replication occurs on template DNA bound by numerous other nucleoprotein complexes involved in other important cellular functions such as transcription, DNA repair, and recombination. However, the presence of these proteins on the template DNA creates potential roadblocks to the replisome, which can impede replication and result in fork collapse and genome instability (9–11). Replication across roadblocks requires the action of accessory proteins or activation of DNA repair pathways and replication restart mechanisms (12).

Both prokaryotes and eukaryotes express accessory replicative helicases that remove protein roadblocks from the path of the replication fork. These enzymes often act on the strand opposite to that encircled by the ring-shaped replicative helicase to rescue stalled replication forks and prevent the collapse of the progressing replisome (12, 13). In eukaryotes, the Pif1 helicase is important for replisome bypass of R-loops and protein roadblocks (14–16). In *E. coli*, the Rep and UvrD helicases promote the removal of a range of roadblocks, including RNA polymerase (RNAP) (17–23). This underlying activity is believed to be dependent on coordination with either a stalled or progressing replisome (14,17,24), but the precise mechanisms are not completely understood.

The *E. coli* Rep helicase is a superfamily 1A (SF1A) helicase that translocates on single-stranded DNA (ssDNA) in a 3ʹ–5ʹ direction (25). Rep like its structural homologs *E. coli* UvrD and *Bacillus stearothermophilus* PcrA (reviewed in (12)) comprises of four domains (1A, 2A, 1B and 2B). While the motor cores, termed 1A and 2A, consist of two highly conserved RecA-like subdomains, it is the 2B subdomain that exhibits extreme conformational changes between a closed and open state that are tightly linked to the activation of helicase activity (12, 25). Although Rep, UvrD, and PcrA monomers can bind and translocate on ssDNA, they do not exhibit processive double-stranded DNA (dsDNA) unwinding activity (26–28). However, intramolecular coupling of the 2B subdomain in the extreme closed state, or deletion of the subdomain altogether, has revealed activation of monomeric helicase activity (12,29,30). Additionally, this subdomain has been proposed to play an important role in separating the primary functions of Rep; protein-roadblock displacement and unwinding of dsDNA (12). The displacement of protein roadblocks has proven essential to cell viability. Notably, in *E. coli* UvrD can also rescue stalled replication forks, where single *rep* and *uvrD* mutations are viable but double *rep*, *uvrD* mutations are lethal (17).

Rep can physically interact with the replication fork through both protein-DNA and protein-protein interactions. The opposite translocation direction to that of the lagging-strand DnaB replicative helicase (5ʹ to 3ʹ) places Rep on the leading-strand DNA template (31). Physical interactions between the C-terminus of Rep and DnaB are important in promoting the efficient removal of protein roadblocks. Recent *in vivo* studies showed a maximal occupancy of the DnaB hexamer, revealing six Rep monomers associated with the replisome (24). Rep proteins with a mutated C-terminus showed a decrease in this occupancy. The authors proposed that the C-terminus plays an important role in recruiting Rep monomers to the replisome by the interaction with DnaB (24). Further, this interaction allows the Rep monomers to be loaded onto the leading strand to translocate ahead of the replication fork (24). This was further supported by study of an ATPase deficient mutant of Rep that could colocalize with the replisome but showed no evidence of translocation away from the replication fork. Additionally, the 2B subdomain of Rep is crucial for the displacement of protein roadblocks (32, 33). Positioned at the leading edge of the helicase, conformational changes of this subdomain likely cause the translocation activity of Rep to switch to protein displacement upon contacting potential roadblocks (32, 34). Despite the extensive structural and functional knowledge of Rep and homologs, the kinetic mechanisms underlying the association with the replisome leading to the displacement of roadblocks and stalled replication rescue remain poorly understood.

Investigations of replication fork stalling have resulted in the development of many tools to mimic protein roadblocks. We have previously developed a highly stable, site-specific roadblock using the nuclease dead Cas9 (dCas9) protein (35). This protein roadblock containing an RNA:DNA hybrid can efficiently block bacterial, viral, and yeast replisomes for long periods (> 20 min). Using this tool, in combination with *in vitro* single-molecule fluorescence imaging, we investigated replisome roadblock bypass with a high degree of spatial and temporal resolution. These single-molecule techniques reveal the heterogeneity of complex biological processes and provide insight into the individual behaviors of single molecules that are not detected by ensemble averaging methods.

To investigate the *E. coli* Rep helicase in the context of elongating and stalled replication forks, we use single-molecule assays to directly visualize the individual Rep proteins at the replisome. Using an *in vitro* reconstituted system and the dCas9 roadblock, we monitor Rep behaviors in real time by imaging fluorescently labeled Rep proteins. We set out to test the hypothesis that Rep is recruited to stalled replication forks, which has been raised by several studies (17,24,36), or whether it might associate with the replisome continually regardless of it being stalled or not. We find that during replication elongation, Rep frequently associates with the replisome in a predominantly monomeric state. Further, in the presence of dCas9 roadblocks, we see efficient removal of the roadblock and rescue of replication. By use of less stable roadblocks, we show that the time elapsed between stalling and rescue of replication is not dependent on the stability of the roadblock, but rather represents a process that occurs after the roadblock has been removed.

## MATERIAL AND METHODS

### Proteins

*Escherichia coli* DNA replication proteins were produced as described previously: the β_2_ sliding clamp (37), SSB (38), the DnaB_6_(DnaC)_6_ helicase-loader complex (referred to as DnaBC) (39), DnaG primase (40), the Pol III τ_3_δδʹχψ clamp loader complex (41), and Pol III αεθ core (41, 42). dCas9-dL5, referred to as dCas9, was produced as described previously (35).

### Overproduction and purification of Rep WT, Rep K28A, and Rep ΔC33

#### Construction of plasmids

To construct plasmid pZX2198 that encodes His_6_-Rep under the control of the T7 promoter, the *rep* gene was amplified by PCR using plasmid pCL771, a derivative of pET3c harboring a *rep* gene, as a template and primers PET3 (5’-CGACTCACTATAGGGAGACCACAAC) and 740 (5’-AAAGAGCTCTTATTTCCCTCGTTTTGCCG) that carry a *Sac*I site immediately downstream of the *rep* gene. The PCR product was cleaned using a Qiagen PCR purification kit, digested with *Nde*I and *Sac*I, and ligated into plasmid pETMCSIII (43) that was pre-digested with *Nde*I and *Sac*I and gel purified. The ligation mixture was transformed into *E. coli* strain AN1459 (43). Colony PCR was then performed to identify colonies harboring the desired plasmid. Plasmid DNAs were extracted and sequences of *rep* were verified by DNA sequencing.

To construct plasmid pZX2199 that encodes His_6_-RepΔC33, a 3ʹ-fragment of the *rep* gene missing the C-terminal 33 residues that are known to interact with the DnaB helicase was amplified by PCR using plasmid pCL771 as template and primers 738 (5’-TACTGGCGAGCTGATCG) and 739 (5’-AAAGAGCTCTTACCAAATCAGATCATCCTG). The PCR product was cleaned and digested with *Mlu*I and *Sca*I and ligated into pZX2198 pre-digested with the same enzymes. The plasmid was then selected and verified as above.

Plasmid pZX2200 encoding His_6_-Rep K28A was constructed by site-directed mutagenesis using the QuikChange protocol (manufacturer) with pZX2198 as template and primers 734 (5’-CGCGGGTTCCGGTGCAACTCGTGTTATCACC) and 735 (5’-GATAACACGAGTTGCACCGGAACCCGCGCC).

#### Protein expression and purification

Proteins were over-expressed using *E. coli* strain BL21(DE3)/pLysS harboring the desired plasmids. Briefly, a bacterial colony was inoculated into LB medium supplemented with 100 µg mL^-1^ of ampicillin and grown at 37°C overnight, then 5 mL of overnight culture was inoculated into 1 L of LB supplemented with 100 µg mL^-1^ of ampicillin (2 L in total). Bacteria were grown at 37°C until OD_600_ of the culture was approximately 0.6. Protein expression was induced by adding IPTG to a final concentration of 0.4 mM. The cultures were then grown for 3 h at 25°C and the cells were collected by centrifugation. Cell pellets were weighed, snap-frozen in liquid nitrogen, and stored at –80°C until use.

Bacterial cells were resuspended in 60 mL of Lysis buffer containing 50 mM Tris-HCl, pH 7.6, 10% sucrose, 300 mM NaCl, 2 mM dithiothreitol, 1 mM EDTA. A tablet of EDTA-free protease inhibitor cocktail (Roche) was also added. The cells were lysed by two passages through a French Press operated at 12,000 psi. Cell debris was removed by centrifugation (40,000 × *g*, 20 min). Ammonium sulfate (0.31 g mL^-1^) was then added to the cleared lysate and stirred for 1 h. Precipitated proteins were collected by centrifugation (40,000 × *g*, 50 min) and dissolved in 50 mL of Buffer A (50 mM Tris-HCl, pH 7.6, 10% glycerol, 700 mM NaCl, 0.5 mM dithiothreitol, 15 mM imidazole). The protein was loaded onto a 5 mL HisTrap HP column (Cytiva) equilibrated with Buffer A. The loaded column was washed with 100 mL of Buffer A and proteins were eluted with a gradient of 15−600 mM imidazole over 25 mL. Fractions containing Rep proteins were pooled and diluted 2.5-fold with Buffer A. The solution was loaded onto a 5 mL HiTrap Heparin column (Cytiva) equilibrated with Buffer B (50 mM Tris-HCl, pH 7.6, 30% glycerol, 1 mM EDTA, 1 mM dithiothreitol, 0.2 M NaCl). The loaded column was washed with 40 mL of Buffer B and proteins were eluted with a gradient of 0.2–1.0 M NaCl over 120 mL. The fractions containing pure Rep proteins were combined and dialyzed against 1 L of a storage buffer containing 50 mM Tris-HCl, pH 7.6, 30% (*v*/*v*) glycerol, 450 mM NaCl, 2 mM dithiothreitol and 1 mM EDTA, and stored at –80°C. Purity of samples was confirmed by SDS-PAGE (Supplementary Figure S1).

### Expression, purification and labeling of Rep (A97C)

Rep (A97C) was expressed and purified as previously described (44), with modifications. Rep (A97C) was overproduced using *E. coli* BL21(DE3) harboring the pRepA97C plasmid (43). Briefly, a bacterial colony was inoculated into 30 mL of LB medium supplemented with 30 µg mL^-1^ of kanamycin in a 100 mL flask and grown at 37°C overnight. 10 mL of overnight culture was inoculated into 1 L of LB supplemented with 30 µg mL^-1^ of kanamycin (2 L in total). Bacteria were grown at 30°C until OD_600_ ∼ 0.8, and protein expression was induced by the addition of 300 µM IPTG. After growth for 2 h at 30°C, cells were collected by centrifugation, weighed, snap-frozen in liquid nitrogen and stored at –80°C.

Thawed cells were resuspended as described above, with the following modifications. Immediately before lysis, 0.5 mM phenylmethylsulfonyl fluoride (PMSF) was added. Following lysis by French Press, cell debris was removed by centrifugation (40,000 × *g*, 30 min). Dissolved, precipitated proteins were purified as described above, where Heparin purification was carried out with Buffer B containing 10% glycerol. Fractions containing pure Rep proteins were combined and dialyzed against 2 L of a storage buffer containing 50 mM Tris-HCl, pH 7.6, 30% (*v*/*v*) glycerol, 300 mM NaCl, 5 mM dithiothreitol, and 1 mM EDTA; and stored at –80°C. The purity of the sample was confirmed by SDS-PAGE (Supplementary Figure S1).

Purified Rep (A97C) was fluorescently labeled with Alexa Fluor 647 (Invitrogen) using methods adapted from (45, 46). First, a total of 1 mg of Rep (A97C) was reduced with 5 mM tris(2-carboxyethyl)phosphine (pH 7.6) in storage buffer containing 50% (*v*/*v*) saturated ammonium sulfate solution, at 6°C for 1 h with gentle rotation to yield Fraction I. Fraction I was centrifuged (21,000 ×*g*, 15 min) at 6°C and the supernatant was carefully removed. The precipitate was washed with ice-cold labeling buffer (50 mM Tris-HCl, pH 7.6, 30% (*v/v*) glycerol, 300 mM NaCl, 1 mM EDTA) and 50% (*v/v*) saturated ammonium sulfate solution to yield Fraction II (both solutions had been extensively degassed by sonication and deoxygenated using Ar gas). Fraction II was incubated at 6°C for 1 h with gentle rotation, then centrifuged (21 000 × *g*, 15 min) at 6°C and the supernatant was removed to yield Fraction III. The labeling reaction was carried out on Fraction III, now devoid of reducing agent, using a 5-fold molar excess of maleimide-conjugated AF647 with 33 µM Rep (A97C) in 300 µL of deoxygenated and degassed labeling buffer. The reaction was allowed to proceed at 6°C overnight with gentle rotation (in the dark), quenched with 30 mM dithiothreitol for 1 h at 6°C to yield Fraction IV. Fraction IV was split into three equal volumes and applied to Zeba Spin desalting columns (40K MWCO) (ThermoFisher) equilibrated with storage buffer. Free dye was separated from labeled protein by centrifugation (1500 × *g*, 2 min) at 6°C. The flow-through from each column was then applied to a second desalting column and centrifuged. The flow-through from each column was combined and stored at –80°C. The degree of labeling was determined by UV/Vis spectroscopy to be 1.0 fluorophore per Rep monomer. Alexa Fluor 647-labeled Rep (A97C) is henceforth referred to as Rep-AF647.

### Surface plasmon resonance

SPR experiments were carried out on a BIAcore T200 instrument (Cytiva) using a streptavidin (SA) coated sensor chip to study the binding and dissociation of Rep WT, Rep K28A, and Rep ΔC33 to ssDNA substrates. All experiments were carried out at 20°C at a flow rate of 10 (chip preparation) or 20 µL min^-1^ (Rep protein binding). The SA chip was activated with four sequential 40 s injections of 1 M NaCl, 50 mM NaOH, then stabilized by 1 min treatment with 1 M MgCl_2_.

The 3ʹ-biotinylated-dT_35_, -dT_15_ or –dT_10_ substrates were dissolved in 1 × SPR buffer (25 mM Tris-HCl, pH 7.6, 50 mM NaCl, 5 mM MgCl_2_, 0.25 mM dithiothreitol, 0.2 mM EDTA and 0.005% (*v/v*) surfactant P20) to a final concentration of 10 nM and introduced onto the SPR chip for immobilization, followed by three sequential wash steps with 1 M MgCl_2_. The signal from the ssDNA substrates corresponded to 250 response units (RU) (dT_35_), 170 RU (dT_15_) and 125 RU (dT_10_).

Following the immobilization of the ssDNA substrates, binding studies were done by injecting specified concentrations of Rep proteins in the SPR buffer. For measurements of His_6_-Rep (WT, K28A, and ΔC33) binding to dT_35_ and dissociation in the presence of 200 µM of AMP-PNP, ADP, or ATP, 20 nM of protein was injected for 60 s, followed by injection of the specified nucleotide cofactor for 60 s. For measurements of His_6_-Rep WT binding to dT_35_, dT_15_, and dT_10_, 20 nM of His_6_-Rep WT was injected for 60 s, followed by injection of SPR buffer lacking any Rep or nucleotide cofactor. For measurements of His_6_-Rep WT binding to dT_15_, an optimized concentration range of [0, 1, 2, 4, 8] µM was injected at 60 µL min^-1^ for 60 s.

### Rolling-circle replication templates

The 2-kbp DNA rolling-circle substrates were prepared as previously described (47). The 18-kbp DNA rolling-circle substrates were prepared from plasmid pUBER using methods and oligonucleotides described by Mueller et al. (48). Briefly, 50 μg of supercoiled pUBER plasmid was treated with 1 unit/μg Nt.*Bbv*CI in 1 × Cutsmart buffer (New England Biolabs, USA) at 37°C for 4 h. A 10-fold molar excess of Dig-competitor oligonucleotides was added to the reaction and the temperature was raised to 65°C for 20 min followed by cooling to 14°C at a rate of 1°C min^-1^. The displaced oligonucleotides were purified from the gapped plasmid by magnetic separation using 1 μg/nmol tosyl activated paramagnetic beads functionalized with anti-digoxigenin Fab fragments, equilibrated with 1× Cutsmart buffer. The nicking reaction mixture was incubated with functionalized beads for 30 min at room temperature with gentle rotation. The fork oligonucleotide was annealed to the gapped plasmid in the presence of 100-fold molar excess over the DNA substrate in 1× Cutsmart buffer at 50°C for 10 min followed by cooling to 16°C at 1°C min^-1^. The fork oligonucleotide was ligated to the DNA substrate by adding 62.5 units/μg T4 DNA ligase, supplementing with 12 mM ATP and 10 mM dithiothreitol, and incubating at 16°C for 18 h followed by inactivation of the ligase at 65°C for 10 min. 18-kbp rolling-circle DNA templates were purified from excess fork oligonucleotides by gel filtration chromatography on a Sepharose 4B (1 × 25 cm; Sigma-Aldrich) column in the presence of 12 mM EDTA and 300 mM NaCl. Rolling-circle DNA templates were eluted with TE buffer containing 10 mM Tris-HCl, pH 7.6, 1 mM EDTA and 300 mM NaCl. Fractions containing the 18-kbp rolling-circle DNA template were stored at – 80°C.

### *In vitro* ensemble replication rescue assays

Standard leading-strand replication assays were set up as described previously (35), with the following modifications. Replication rescue assays were set up in replication buffer (RB; 30 mM Tris-HCl pH 7.5, 12 mM magnesium acetate, 50 mM potassium glutamate, 0.5 mM EDTA and 0.0025% (*v/v*) Tween20) or RB containing high magnesium (RBM; containing 24 mM magnesium acetate). Reactions contained 2 nM 2-kbp rolling-circle replication template, specified concentrations of dCas9 and gRNA, 60 nM DnaBC, 30 nM τ_3_δδʹχψ, 90 nM Pol III αεθ core, 200 nM β_2_, 10 mM dithiothreitol, 1 mM ATP (in RB) or 10 mM ATP (in RBM), and 125 µM dNTPs (each) in a final volume of 12 µL. First, dCas9 was incubated with the specified cgRNA for 5 min at room temperature, and further incubated with rolling-circle DNA templates for 5 min at room temperature. Other components were mixed and incubated at room temperature, then cooled on ice for 5 min before addition of dCas9-cgRNA-DNA complexes. Reactions were initiated at 30°C. At specified time points, stated concentrations of His_6_-Rep WT, His_6_-Rep K28A, His_6_-Rep ΔC33, or Rep-AF647 were added to the reactions in the absence or presence of 50 nM trap dsDNA, an 83-mer target DNA containing one complementary sequence to the fully complementary gRNA (cgRNA1) (Supplementary Table S1)(35). The reactions were quenched at specified time points by the addition of 12 μL of LES (2 × DNA gel loading dye, 200 mM EDTA and 2% (*w*/*v*) SDS). The quenched reactions were loaded into a 0.6% (*w/v*) agarose gel in 2 × TAE. Products were separated by agarose gel electrophoresis at 60 V for 150 min, stained in SYBR-Gold (Invitrogen), and imaged under long-wave UV light.

*E. coli* leading- and lagging-strand DNA replication reactions were set up as previously described (35, 42), with minor modifications. Reactions were set up in RB and contained 4 nM 2-kbp rolling-circle DNA template, specified concentrations of dCas9 and cgRNA, 60 nM DnaBC, 80 nM DnaG, 30 nM τ_3_δδʹχψ, 10 nM SSB, 90 nM Pol III αεθ core, 200 nM β_2_, 10 mM dithiothreitol, 1 mM ATP, 125 µM dNTPs and 250 µM NTPs to a final volume of 12 µL. At 10 min after initiation of the replication reaction, a specified concentration of His_6_-Rep WT was added to each reaction in the presence of 50 nM trap dsDNA. At 20 min, reactions were quenched by the addition of 1.5 µL 0.5 M EDTA and 3 µL DNA loading dye (6 mM EDTA, 300 mM NaOH, 0.25% (*w*/*v*) bromocresol green, 0.25% (*w*/*v*) xylene cyanol FF, 30% (*v*/*v*) glycerol). DNA products were separated on a 0.6% (*w*/*v*) alkaline agarose gel at 14 V for 14 h. The gel was neutralized in TAE buffer, stained with SYBR-Gold and imaged under UV light.

### *In vitro* single-molecule fluorescence microscopy

*In vitro* single-molecule fluorescence microscopy was carried out on an Eclipse Ti-E inverted microscope (Nikon, Japan) with a CFI Apo TIRF 100× oil-immersion TIRF objective (NA 1.49, Nikon, Japan) as described previously (35,39,42,46). The temperature was maintained at 31.2°C by an electronically heated flow-cell chamber coupled to an objective heating jacket (Okolab, USA). NIS-elements software was used to operate the microscope and the focus was locked into place through the Perfect Focus System (Nikon, Japan). Images were captured using a 512 × 512 pixel EM-CCD camera (either Photometrics Evolve 512 Delta or Andor iXon 897). DNA molecules stained with 150 nM Sytox orange were imaged with either a 568-nm laser (Coherent, Sapphire 568-200 CW) at 400 mW cm^-2^, a 514-nm laser (Coherent, Sapphire 514-150 CW) at 200 mW cm^-2^, or 532-nm laser (Coherent, Sapphire 532-300 CW) at 90 mW cm^-2^. dCas9 complexed to cgRNA-Atto647 was imaged with a 647-nm laser (Coherent, Obis 647-100 CW) at 220 mW cm^-2^. Rep-AF647 was imaged with the 647-nm laser at 200 mW cm^-2^.

#### Preparation of flow cells for in vitro imaging

Replication reactions were carried out in microfluidic flow cells constructed from a PDMS flow chamber placed on top of a PEG-biotin-functionalized glass microscope coverslip as described previously (41,42,49,50). Once assembled, all surfaces of the flow cell including tubing were blocked against non-specific binding by the introduction of at least 300 µL blocking buffer (50 mM Tris-HCl pH 7.9, 50 mM potassium chloride, 2% (*v/v*) Tween20).

#### Single-molecule Rep binding assay

The single-molecule Rep binding assay was designed to investigate the association of Rep to forked DNA templates bound by DnaBC and SSB, in the absence and presence of ATP. First, 8 pM 2-kbp rolling-circle DNA template was incubated with 7 nM DnaBC in degassed single-molecule replication buffer (SM; 25 mM Tris-HCl pH 7.9, 10 mM magnesium acetate, 50 mM potassium glutamate, 0.1 mM EDTA, 0.0025% Tween20, 0.5 mg mL^-1^ BSA, 1 mM ATP, 10 mM dithiothreitol and 150 nM Sytox orange) for 3 min at 37°C. Following, 20 nM SSB was added to the DNA-DnaBC mixture. The DNA-DnaBC-SSB was adsorbed to the flow-cell surface at 10 µL min^-1^ until an appropriate surface density was achieved. The flow cell was subsequently washed with SM buffer containing Sytox orange at 70 µL min^-1^ for 2 min. Next, 10 nM Rep-AF647 in SM buffer supplemented with 10 nM SSB, 10 mM dithiothreitol, and 150 nM Sytox orange, in the presence or absence of 5 mM ATP, was added to the flow cell at 50 µL min^-1^ for 1 min and then at 10 µL min^-1^ for 5 min. To detect Rep binding to the DNA, the Sytox orange-stained rolling-circle DNA template was visualized with a 532-nm laser (90 mW cm^-2^) sequentially for 200 ms once every second for 3 min. The Rep-AF647 protein was visualized with a 647-nm laser (200 mW cm^-2^) sequentially for 200 ms once every second for 3 min. Fluorescence signals were captured with an EMCCD camera (Andor iXon 897) with appropriate filter sets.

#### Single-molecule rolling-circle replication assay

The single-molecule rolling-circle replication assays were carried out in microfluidic flow-cell devices with the same DnaBC pre-incubation step and DNA adsorption step as described above. The replication step was carried out as previously described (35,39,42,46) with modifications, described below.

#### Visualization of Rep during replication

The replication solution contained specified concentrations of His6-Rep WT or Rep-AF647, 30 nM Pol III αεθ core, 10 nM τ_3_δδʹχψ, 46 nM β_2_, 75 nM DnaG and 20 nM SSB_4_ in SM buffer plus 250 µM of each NTP, 50 µM of each dNTP. Replication was initiated by injecting the replication solution into the flow cell containing immobilized DNA-DnaBC at 70 µL min^-1^ for 1 min and then slowed to 10 µL min^-1^ for 10 min. Sytox orange stained DNA molecules were imaged with a 514-nm laser (200 mW cm^-2^) sequentially for 200 ms once every second for a period of 1 min. The Rep-AF647 protein was visualized with a 647-nm laser (200 mW cm^-2^) sequentially for 200 ms once every second for 1 min.

#### Replication rescue at pre-incubated roadblocks

Visualization of replication rescue by His_6_-Rep WT, His_6_-Rep K28A, or His_6_-Rep ΔC33 in conditions where the DNA template has been pre-incubated with the dCas9 roadblock were carried out as described previously (35) with the following modifications. 5 nM dCas9 was incubated with 20 nM cgRNA1-Atto647 for 5 min at 37°C in SM buffer (omit Sytox orange). The dCas9-cgRNA1-(Atto647) complex, referred to as dCas9-cgRNA1-Atto647, was then incubated with 80 pM 2-kbp rolling-circle DNA template for 5 min at 37°C. To form the DNA pre-incubation solution, the DNA-dCas9-cgRNA1-Atto647 complex was then incubated with 70 nM DnaBC for 3 min at 37°C. The DNA pre-incubation solution was then diluted 1:10 with SM buffer plus 150 nM Sytox orange and adsorbed to the surface of the flow cell at 10 µL min^-1^ until an appropriate DNA density was achieved. The flow cell was then washed with 200 µL of SM buffer. Following this, replication was initiated with the continuous presence of replication proteins as above in the absence or presence of 20 nM His_6_-Rep WT, His_6_-Rep K28A, or His_6_-Rep ΔC33. DNA and dCas9-cgRNA1-Atto647 were imaged in multiple fields of view per experiment, by sequential excitation with a 532-nm laser (90 mW cm^-2^) and a 647-nm laser (200 mW cm^-2^) for 200 ms once every 10 s for 3 min.

#### Replication rescue with roadblocks in solution

Visualization of replication rescue by Rep-AF647 in conditions where both Rep and dCas9-cgRNA complexes were in solution with the replisome components was carried out as previously described (35,39,42,46), with the following modifications. First, the dCas9-cgRNA complex was formed with the specified cgRNA as described for the pre-incubation of roadblocks with DNA assay. Next, the DNA-DnaBC complex was formed and adsorbed to the flow-cell surface as described previously. Following, the replication solution was mixed as described for visualization of Rep during elongating replication assays, with the addition of specified concentrations of dCas9-cgRNA (determined by dCas9 concentration in the complex) and Rep-AF647. Reactions were initiated with the addition of the replication solution to the flow cell at 70 µL min^-1^ for 1 min and then slowed to 10 µL min^-1^ for 10 min. DNA and Rep were imaged in one field of view per experiment, by sequential excitation with a 532-nm laser (90 mW cm^-2^) and a 647-nm laser (200 mW cm^-2^) for 200 ms once every second for 4 min.

#### Single-molecule characterization of mismatch gRNA binding durations on short oligos

To assess the duration of binding of mismatch gRNAs (MMgRNA) in complex with dCas9, an 83-bp oligo containing one complementary sequence to the fully complementary gRNA (cgRNA1) was used (Supplementary Table S1) (35). First, the dCas9-MMgRNA complexes were formed by pre-incubating 5 nM dCas9 with 20 nM of the specified MMgRNA-Atto647 in SM buffer (with ATP omitted throughout) for 5 min at 37°C. Next, 1 pM of 83-bp oligo was adsorbed to the flow-cell surface in SM buffer at 10 µL min^-1^ for 2 min. The dCas9 pre-incubation mixture was then diluted 1:10 with SM buffer and flowed into the flow cell at 70 µL min^-1^ for 1 min. The buffer was then switched immediately to only SM buffer and flowed at 10 µL min^-1^ for 20 min. The DNA oligos and dCas9-MMgRNA-Atto647 complexes were imaged in multiple fields of view per experiment, by sequentially exciting with a 532-nm laser (140 mW cm^-2^) and a 647-nm laser (200 mW cm^-2^) for 200 ms once every 30 s for 10 min.

### Data Analysis

All analyses were carried out using ImageJ/Fiji (1.51w), MATLAB 2016b, OriginPro 2021b and in-house built plugins, found here https://doi.org/10.5281/zenodo.7379064.

#### Degree of labeling

The number of fluorophores per labeled Rep-AF647 was quantified at the single-molecule level by immobilizing 40 pM Rep-AF647 onto the flow cell surface in SM buffer (less ATP). The fluorophores were imaged by exciting for 200 ms constantly for 3 min until the fluorophores were photobleached. Raw movies were corrected for the electronic offset of the camera and excitation-beam profile. Single molecules of Rep-AF647 were identified using an in-house built peak fitter tool and the photobleaching steps were fit using change-point analysis (51–53). The histogram of steps per molecule showed a degree of labeling of mostly 1 dye per Rep monomer (Supplementary Figure S2). It is likely observations of more than 2 fluorophores represent more than a monomer in the identified peak observed, as there is only one cysteine residue in each monomer.

#### Automatic analysis of rolling-circle DNA replication

To automatically track the replication of the rolling-circle DNA templates, raw videos (.nd2 format) were first converted to TIF files and flattened with the excitation beam profile, as previously described (54). Detectable drift between subsequent frames was then corrected and any un-replicated molecules were removed by subtracting the first frame from each consecutive frame. This prevents un-replicated DNA templates on the surface from interfering with the detection and tracking of replicating molecules. Next, replicating molecules of interest were selected and analyzed individually. Positions of the replicating molecule were determined with custom-written ImageJ and MATLAB plug-ins that detect the leading edge of each replicating molecule, saving as coordinates for downstream analysis. These coordinates were used to detect individual rate segments of the replicating molecules by change-point analysis.

#### Automatic tracking of labeled Rep during rolling-circle DNA replication

Videos of labeled Rep proteins during rolling-circle DNA replication were flattened and prepared as above. Using the coordinates saved from automatic tracking of the replicating DNA molecule, Rep-AF647 molecules were detected within the proximity of the tip of the replicating molecule by using custom-written ImageJ and MATLAB plug-ins. Here, regions of interest around the replicating tip of the DNA molecules were expanded and transposed into the corresponding video of Rep-AF647. Peak finder was used to detect any Rep-AF647 molecules and colocalization was determined for those peaks that were within the distance of the tip of the replicating DNA molecule. The intensities of colocalized Rep-AF647 molecules were measured and individual binding events were detected by change-point analysis.

#### Determination of stoichiometry of Rep

The number of labeled Rep molecules binding to DNA, at actively replicating or stalled replisomes, was calculated by dividing their initial intensities by the intensity of a single fluorophore, as previously described (39, 42). Briefly, the average intensity per fluorophore was quantified by detecting photobleaching steps in labeled Rep non-specifically bound to the coverslip surface (Supplementary Fig S2). Imaging was done under the same conditions as the experiment of interest. The integrated intensity for every fluorescently labeled Rep visible in the field of view was calculated after applying a local background subtraction. The histograms obtained were fit with a Gaussian distribution function to give the average intensity of a single molecule.

#### Colocalization analysis

Sytox orange stained DNA molecules and dCas9-cgRNA-Atto647 complexes, dCas9-MMgRNA-Atto647, or Rep-AF647 molecules were classed as being colocalized as previously described (39). Briefly, foci of interest were classed as being colocalized if their centroid positions (determined using an ImageJ in-house built peak finder tool) fell within 2 pixels of each other. The chance of coincidental colocalization (*C*) was calculated using Equation 1, where *A*_R_ is the focus area, *A*_FOV_ is the field of view area, and *n* is the number of foci.

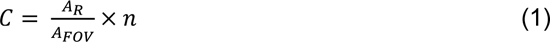

#### Determination of Rep association

Rep association dynamics were extracted by tracking the fluorescence intensity of Rep-AF647 molecules over time. A threshold was applied for each trajectory equivalent to the intensity of half a Rep-AF647 molecule. The binding frequency of Rep-AF647 during replication, was defined as the number of times per minute where the intensity exceeded the threshold.

## RESULTS

### Purified and labeled Rep binds to DNA

Association of Rep, and other similar SF1B helicases, to various DNA structures has been widely characterized by previous studies (reviewed in (55, 56)). We used surface plasmon resonance (SPR) to assess DNA binding and ATP hydrolytic properties of the purified His_6_-Rep variants. First, we assayed the binding of His_6_-Rep WT (400 nM) to 3ʹ-biotinylated-dT_35_ (3ʹ-bio-dT_35_) immobilized onto the surface of a streptavidin-coated SPR chip (Supplementary Figure S3A). We found that binding and dissociation of Rep to and from dT_35_ was biphasic, one phase with faster association and slower dissociation, and the other with slow association and faster dissociation (Supplementary Figure S3A). On the basis of the different dissociation times, we reasoned that the former likely represents Rep dimer binding, and the latter, Rep monomer binding. Additionally, based on the signal amplitude we calculated the equivalent of four Rep monomers binding to the dT_35_. The footprint of a Rep monomer is approximately 8 nt (57), thus 35 nt ssDNA could accommodate 4 Rep monomers resulting in the coating of the ssDNA at high Rep concentrations introduced over long periods.

Introduction of His_6_-Rep WT at a lower concentration (20 nM) resulted in the binding of predominantly Rep dimers to dT_35_, with faster association and slower dissociation (Figure 1B, Supplementary Figure S3B). Binding of Rep to dT_35_ resulted in an association rate of 1.03 ± 0.00 × 10^6^ M^-1^s^-1^ and dissociation rate of 2.15 ± 0.03 × 10^-4^ s^-1^ Local fitting of association and dissociation with mass transfer provided an estimated *K*_D_ for Rep dimer:ssDNA of 209 ± 1 pM. Further, we interrogated the binding of His_6_-Rep WT to shorter ssDNA oligonucleotides (dT_10_ and dT_15_). Here, we observed the binding of Rep to shorter ssDNA equivalent to monomers with an association rate of 1.22 ± 0.00 × 10^7^ M^-1^s^-1^. Bound Rep proteins dissociated much faster than Rep bound to dT_35_, with dissociation rates estimated to be 5.91 ± 0.02 × 10^-3^ s^-1^ for dT_15_ (Figure 1B). To estimate the *K*_D_ of the Rep monomer binding to dT_15_, we titrated His_6_-Rep WT at different concentrations (1, 2, 4, and 8 nM) (Supplementary Figure S3C). Global fitting of association and dissociation (1:1 binding with mass transfer) yielded a *K*_D_ for Rep monomer:ssDNA of 484 ± 2 pM. Thus, we hypothesize that dT_10_ and dT_15_ cannot accommodate the predicted 16-nt footprint of the Rep dimer, which we show is the stable form of Rep on ssDNA with a 2.5-fold lower *K*_D_.

**Figure 1.**
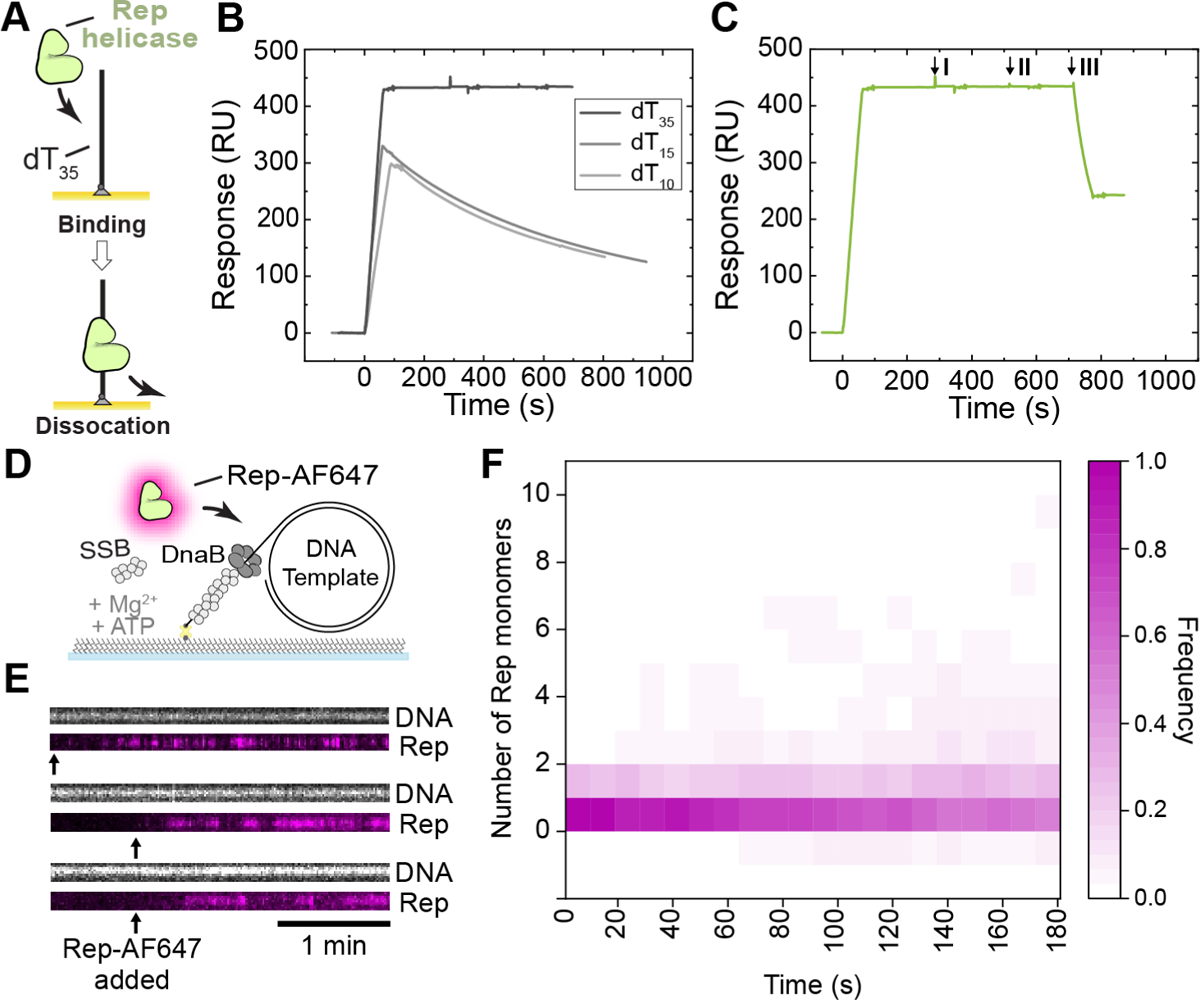
Visualization of Rep binding to ssDNA. **(A)** Schematic representation of Rep proteins binding to dT_35_ oligos in surface plasmon resonance investigations. (B–C) SPR sensorgrams of **(B)** 20 nM Rep WT association (60 s) and dissociation from dT_35_ (dark gray), dT_15_ (gray) and dT_10_ (light gray), and **(C)** Rep WT response to interrogation of 200 μM AMP-PNP (I), ADP (II), and ATP (III) injection at 20 μL min^-1^ for 60s. Only ATP resulted in fast dissociation of Rep from dT_35_. **(D)** Schematic representation of the single-molecule Rep-AF647-DNA binding assay. Rolling-circle DNA templates (2030 bp) are pre-incubated with DnaBC and applied to a microfluidic flow cell. The 5ʹ-biotinylated DNA template couples to the streptavidin functionalized coverslip. Rep-AF647, in the presence of SSB, ATP, and magnesium, is then applied to the flow cell and imaged to monitor for binding. **(E)** Three example kymographs of Rep-AF647 (magenta – bottom) binding to DNA templates (gray – top) in the presence of 5 mM ATP. Arrows indicate the time point of addition to the flow cell. **(F)** Heatmap of the number of Rep-AF647 monomers bound to the DNA template over time in the presence of ATP (*n* = 65).

Further, we looked at the effect of nucleotides on the affinity of His_6_-Rep WT, K28A, and ΔC33 for dT_35_. We show that ATP, but not ADP or AMP-PNP, stimulated the dissociation of His_6_-Rep WT and His_6_-Rep ΔC33 from ssDNA (Figure 1C and Supplementary Figure S3D). Dissociation of His_6_-Rep K28A, cannot be stimulated by ATP (Supplementary Figure S3D). This mutant does not bind or hydrolyze ATP (58), suggesting that ATP hydrolysis is required for the induced faster dissociation of Rep variants from ssDNA. Additionally, ssDNA-binding and ATP-stimulated dissociation of His_6_-Rep WT was similar with dT_35_ immobilized via biotin attached to either the 3’ or 5’ end (3ʹ-bio-dT_35_ and 5ʹ-biotinylated-dT_35_; data not shown), suggesting Rep directly dissociated from DNA upon ATP binding or hydrolysis rather than translocating off the 5ʹ-end. Our measurements are consistent with previous data indicating that the presence of the N-terminal His_6_ tag does not disrupt the DNA binding and ATP hydrolytic properties of Rep (56, 57). Therefore, His_6_-tagged Rep variants, which are easier to purify and free from DNA contamination were used in subsequent studies, and are herein referred to as Rep WT, Rep K28A, and Rep ΔC33.

To visualize Rep behaviors in single-molecule fluorescence assays, we used a mutant of Rep (A97C) (44) site-specifically labeled with a cysteine-reactive red fluorescent dye (Alexa Fluor 647) (referred to as Rep-AF647). Wild-type Rep contains five native and non-essential cysteine residues, all of which have been mutated, and a site-specific cysteine residue replacing Ala97. This residue is within the 1B subdomain and predicted to be close to the 3ʹ-end of the ssDNA (44). This subdomain is not responsible for ATP binding or hydrolysis, or rotation of the 2B subdomain; activities previously characterized to be important for Rep functions (24,30,32–34,59). Additionally, this mutant has been previously used in bulk biochemical and smFRET assays, exhibiting intact helicase and ATPase activities (44).

To characterize Rep-AF647 binding to DNA in single-molecule fluorescence assays we used a 5ʹ-biotin-tailed, gapped and circular DNA template (2030 bp) with replication-fork topology (47). We set out to characterize the binding of Rep-AF647 in the presence and absence of ATP to DNA occupied by DnaBC and SSB (Figure 1D). DnaBC and SSB were pre-incubated with the DNA template and then injected into a microfluidic flow cell to immobilize the complex on a streptavidin-functionalized surface. Rep-AF647, in the presence or absence of ATP, was subsequently injected into the microfluidic flow-cell, and binding events were visualized by near-total internal reflection fluorescence (TIRF) imaging of Sytox orange stained DNA and AF647-labeled Rep molecules. Colocalization and stoichiometry analysis of corresponding foci confirmed that in the absence of ATP, Rep-AF647 binds stably to the DNA template (Supplementary Figure S3E and F). Additionally, significant numbers of Rep-AF647 single-molecules remained bound; equivalent to an average of 8 molecules. This observation suggests that ssDNA in the gapped circular DNA (64 nt) is occupied by a Rep molecule by the end of the acquisition. Therefore, this occupancy in the absence of ATP could suggest that Rep out-competes or interacts with SSB bound to the ssDNA regions. Similar to our observations in the SPR studies, in the presence of ATP this stable binding activity is significantly reduced (Figures 1E and F). Rep-AF647 appears to bind only transiently to DNA in the presence of ATP, suggesting that ATP binding and hydrolysis play an important role in Rep-ssDNA affinity. In agreement with earlier studies (60), we show that ATP binding, and potentially hydrolysis, put Rep into a low-affinity state, resulting in dissociation from the ssDNA template. These data show that labeled Rep protein is active in DNA binding, ATP binding and ATP hydrolysis. Further, our SPR and single-molecule studies confirm the stability of Rep in the absence of ATP on oligonucleotides containing more than 16 nt (56, 57). Together, our results confirm previous hypotheses about Rep-ssDNA affinity and activity.

### Rep associates frequently with elongating replication forks in the absence of roadblocks

Early studies of the growth characteristics of *rep* mutant strains suggested higher replication rates in the presence of Rep (61). Additionally, previous studies have shown that Rep interacts with the DnaB helicase through its C-terminal region (17,58,62). Therefore, we first set out to investigate the effect of Rep WT on the rate and processivity of replication using a single-molecule rolling-circle DNA replication assay. This assay utilizes the 2-kbp rolling-circle template used above, where the 5ʹ-tail is anchored to the surface of the flow cell (7,47,49). Replication is initiated by the introduction of a laminar flow of buffer containing the *E. coli* replication proteins required for coupled leading- and lagging-strand synthesis to form functional replisomes at the fork structure. Initiation of unwinding and synthesis at the fork results in the newly synthesized leading strand being displaced from the circle to form the template for the lagging strand. This process results in the production of a dsDNA tail that is stretched out in the buffer flow and the movement of the circle away from the anchor point, at a rate determined by the replication rate (Figure 2A). These replication events are visualized in real-time by TIRF imaging of Sytox orange-stained dsDNA.

**Figure 2.**
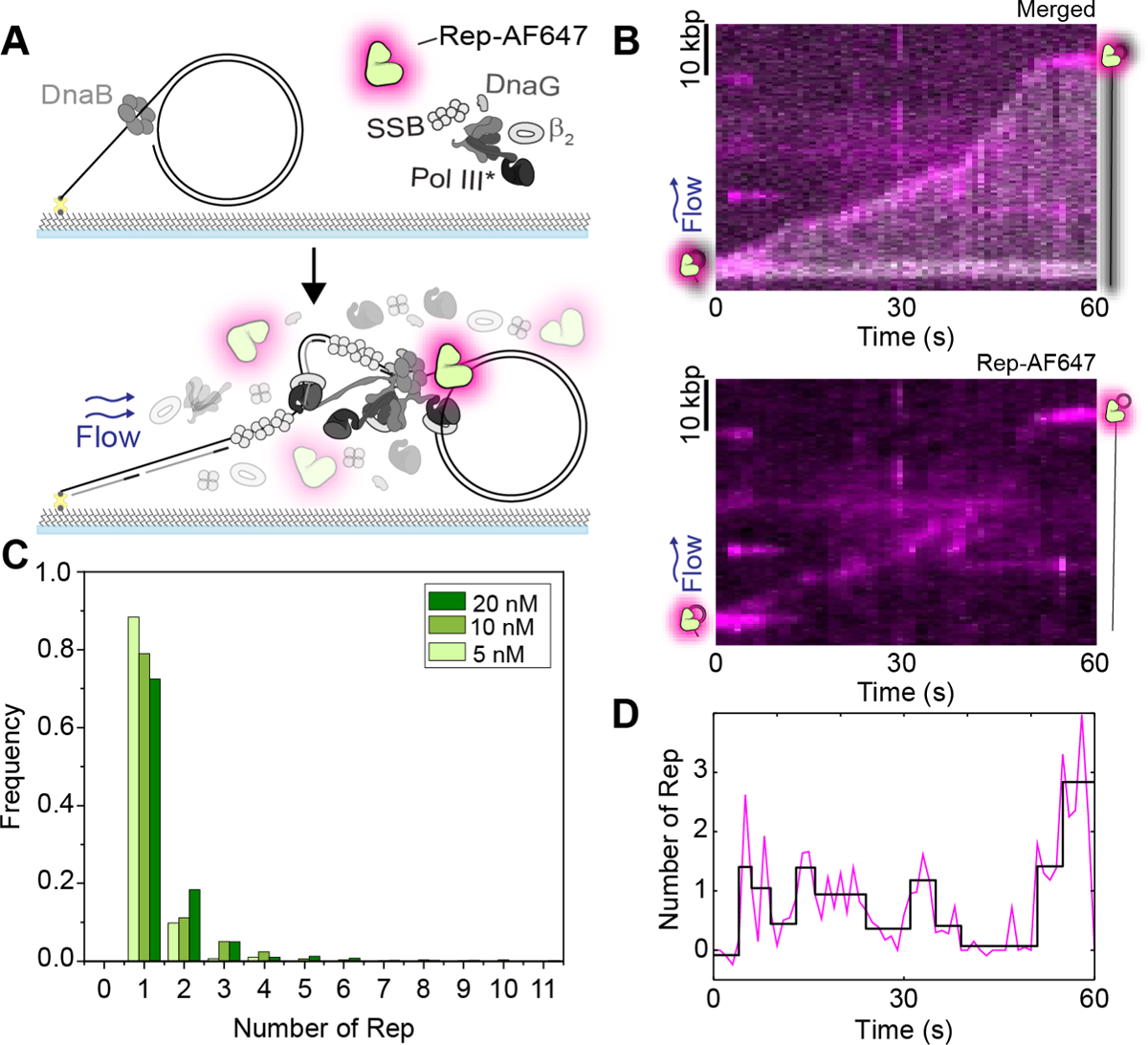
Rep interacts with processive replisomes. **(A)** Schematic representation of the single-molecule rolling-circle DNA replication assay in the presence of Rep-AF647. Rolling-circle DNA templates are pre-incubated with DnaBC and immobilized to the flow cell surface. The addition of the replisome components, ATP, NTPs, and dNTPs, results in the initiation of DNA replication. Sytox orange stained DNA replication products are stretched out by hydrodynamic flow and visualized simultaneously with fluorescent Rep-AF647 proteins. **(B)** Example kymograph of 20 nM Rep-AF647 interacting with the tip of the Sytox orange stained DNA product (merged – top). The Rep-AF647 intensity alone (bottom) shows its frequent association with and dissociation from the replication fork. **(C)** Histogram distributions of Rep-AF647 stoichiometry at the replication fork reveal monomeric stoichiometry at 20 nM (dark green; *n* = 755), 10 nM (green; *n*= 490) and 5 nM (light green; *n* = 286). **(D)** Number of Rep-AF647 as a function of time for the example kymograph in (B) showing variations in Rep stoichiometry at the replication fork during processive replication. Individual steps are detected by change-point analysis (black line).

Analysis of the replication rates of individual replisomes in the absence and presence of unlabeled, wild-type Rep (Rep WT) (5 nM) resulted in median rates of 576 ± 40 and 573 ± 40 bp s^-1^ (median ± SEM), respectively (Supplementary Figure S4A). These rates are in agreement with previously reported *E. coli* replication rates (7,42,46,49). Further, at 10- and 100-fold higher concentrations of Rep WT, resulting replication events revealed rates of 498 ± 33 and 477 ± 40 bp s^-1^, respectively. Taken together, no significant difference was found between median replication rates, suggesting that Rep WT has no effect on the rate of replication at the concentrations used. Further, we saw no significant effect of Rep on the processivity of the replisome, as measured by the length of individual rate segments (Supplementary Figure S4B).

The absence of a clear effect of Rep on DNA replication rate and processivity raises the question of whether Rep is present at all at elongating forks in the absence of protein roadblocks. Previous studies have hypothesized that Rep is only present at replication forks in the event a roadblock is encountered, being that it is recruited to the replication fork upon stalling (17,24,36). To investigate whether Rep is present at elongating replisomes, we added Rep-AF647 to the single-molecule rolling-circle DNA replication assays described above (Figure 2A). This assay provides additional information as it allows simultaneous imaging of fluorescently labeled replisome components and interacting partners during the replication reaction (42,46,63). The addition of Rep-AF647 at a concentration of 20 nM in this assay shows that Rep-AF647 is located at the tip of the growing DNA molecule consistent with its interaction with the elongating replisome (Figure 2B and Supplementary Figure S5). Only 3% of DNA molecules analyzed showed no Rep-AF647 intensity above the threshold at a concentration of 20 nM Rep-AF647. Further, the intensity of interacting Rep-AF647 with the replisome fluctuates throughout the elongation of the DNA product (Figure 2D). A single-exponential fit of binding event durations, as detected by change-point analysis of Rep fluorescence over time, revealed an approximate binding lifetime of 2.0 ± 0.2 s. If the same Rep-AF647 molecule remained bound to the active replisome, the intensity should decay at the characteristic lifetime of photobleaching (8.0 ± 0.1 s) (Supplementary Figure S2E). Therefore, it is likely that Rep-AF647 associates with the replication fork and quickly dissociates to be replaced by a new Rep-AF647 molecule from solution, as observed for Rep-AF647 binding investigations above.

Previous studies have hypothesized the potential of six Rep monomers binding to the replication fork through the hexameric structure of the DnaB helicase (17, 24). Our assays reveal an approximate stoichiometry of 1–2 Rep at the fork at any given time point, independent of the concentration of Rep-AF647 used (5, 10, or 20 nM) (Figure 2C). We determined the average binding frequency of a Rep-AF647 molecule to an elongating replisome by applying thresholding analysis to the Rep-AF647 signal to detect binding events. We found that the average binding frequency during replication elongation was 16 ± 1 min^-1^ at 20 nM Rep-AF647. This average frequency decreased with decreasing Rep-AF647 concentration, to 10 ± 1 and 7 ± 1 min^-1^ at 10 and 5 nM, respectively. Thus, the higher the local concentration of Rep is, the more frequently it associates with the fork. Given the low stoichiometry and regular binding events, these observations suggest potentially two behaviors: 1) that Rep stochastically binds to the replisome, and 2) Rep is interacting with the DnaB helicase, but not occupying all potential binding sites on the hexamer at a given time. Nonetheless, we show that Rep associates with elongating replication forks in the absence of protein roadblocks.

### Wild-type Rep removes model roadblocks and rescues stalled replication forks

To investigate if Rep could remove nucleoprotein complexes and rescue stalled replication forks, we used dCas9 in complex with a complementary guide RNA (dCas9-cgRNA) as a protein barrier. We have previously shown we can block the reconstituted *E. coli* replisome in *in vitro* bulk and single-molecule replication assays using dCas9-cgRNA complexes targeted to a specific site in the rolling-circle template (35). We first tested the ability of Rep WT to remove dCas9-cgRNA complexes targeted to either the leading or lagging strand, in bulk biochemical assays. Here dCas9-cgRNA complexes are incubated with the rolling-circle DNA template, to which the *E. coli* replisome proteins are added. After 10 min of the replication reaction, the indicated concentration of Rep WT is added to the reaction and allowed to proceed for a further 10 min. Finally, the reactions are quenched and products are analyzed by gel electrophoresis.

We show that in reactions containing Rep WT, replication of DNA templates occurs past the dCas9-cgRNA binding site indicating that the dCas9-cgRNA roadblock has been removed (Supplementary Figure S6). This activity occurs in reactions where either the leading or lagging strand is targeted by the dCas9-cgRNA complex (Supplementary Figure S6A). The addition of a trap dsDNA acts to bind free dCas9-cgRNA1 complexes in solution, and thus prevents free dCas9-cgRNA1 complexes from rebinding to the template DNA, where in its absence a laddering in the gel is seen consistent with multiple replication blocking and rescuing events (Supplementary Figure S6B, C). At all concentrations tested in bulk biochemical assays (2–300 nM Rep WT) each reaction resulted in long DNA products showing that Rep has effectively removed the dCas9-cgRNA1 complex allowing replication to proceed (Supplementary Figure S6B, C). Additionally, no replication rescue was observed in reactions containing either the Rep ΔC33 or K28A mutants (Supplementary Figure S6D).

To assess this activity at the single-molecule level, we repeated the above experiments by pre-incubating DNA templates with the dCas9-cgRNA1-Atto647 complex. This allowed us to simultaneously visualize both the potential loss of the dCas9-cgRNA1-Atto647 complexes and the growing replication products, by collecting the Atto647 fluorescence emission and dsDNA-bound Sytox orange fluorescence emission, respectively. Pre-incubation of the DNA template with the roadblock complex allows replication to proceed until the block is encountered (Figure 3A). Before adsorption to the flow-cell surface, the DNA-dCas9-cgRNA1-Atto647 complex is pre-incubated with the DnaBC complex.

**Figure 3.**
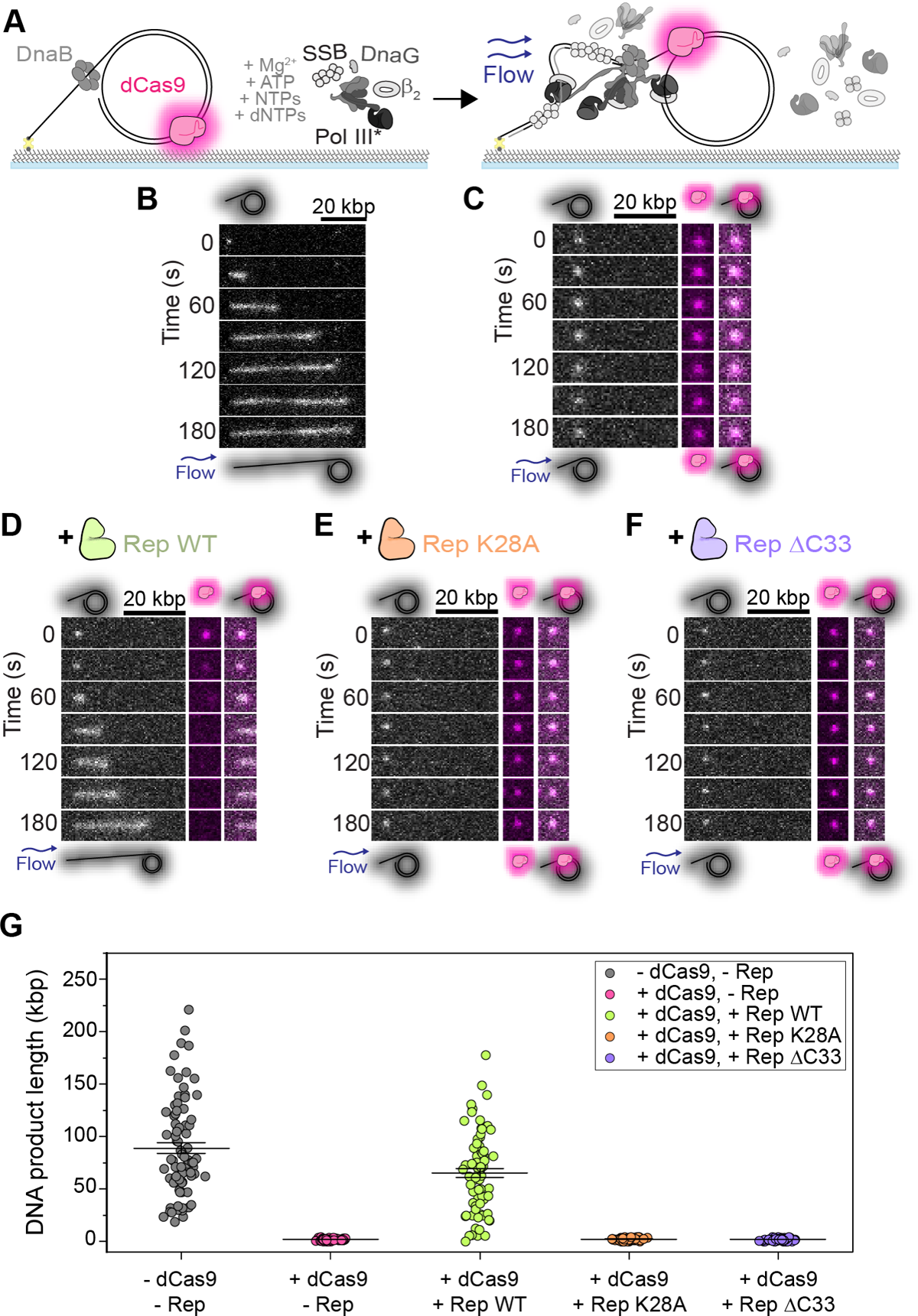
Visualization of stalled replication rescue by Rep at the single-molecule level. **(A)** Schematic representation of single-molecule stalled rolling-circle replication assay. The dCas9-cgRNA1-Atto647 complex is pre-incubated with the rolling-circle DNA template, and further incubated with DnaBC before immobilization to the flow cell. The addition of the replisome components results in the initiation of replication until the dCas9-cgRNA1-Atto647 roadblock has been encountered. Stalled replication is imaged by visualizing the Sytox orange stained-DNA (gray) and dCas9-cgRNA1-Atto647 complex (magenta). (B-F) Example montages and (G) mean DNA product length of **(B)** rolling-circle DNA replication in the absence of protein roadblocks and Rep proteins (89 ± 5 kbp (*n* = 81; replication efficiency of 4.0 ± 0.3% (S.E.M.)), **(C)** stalled DNA replication by dCas9-cgRNA1-Atto647 complex in the presence of all replisome components (2.0 ± 0.1 kbp (*n* = 80; no replicating products observed)), **(D)** stalled replication rescue by Rep WT following removal of the dCas9-cgRNA1-Atto647 complex (65 ± 4 kbp (*n* = 75; 1.6 ± 0.1%)), **(E)** stalled replication in the presence of Rep K28A (2 ± 1 kbp (*n* = 80; no replicating products observed)), and **(F)** stalled replication in the presence of Rep ΔC33 (2 ± 1 kbp (*n* = 80; no replicating products observed)).

Consistent with the bulk assays and previous studies, we saw effective replication blocking of the *E. coli* replisome in the presence of the dCas9-cgRNA1-Atto647 complex (Figure 3B, C, and G, Supplementary Figure S7), indicated by the synthesis of short replication products (35). In the presence of Rep WT, long DNA products are synthesized following the removal of the dCas9-cgRNA1-Atto647 complex (Figure 3D and G, Supplementary Figure S7). Observations of dCas9-cgRNA1-Atto647 complexes photobleaching when imaged every 200 ms, showed a characteristic photobleaching lifetime of 87 s (Supplementary Figure S7D). However, in these experiments, the dCas9-cgRNA1-Atto647 complexes are only imaged once every 10 s, extending the lifetime of the fluorophore. Thus, we are confident that the loss of the dCas9-cgRNA1-Atto647 intensity is due to the displacement of the complex by Rep WT. Additionally, unlike the ensemble assays, displaced dCas9-cgRNA1-Atto647 complexes are carried away from replicating templates in the buffer flow, thus preventing rebinding to the template. In the presence of the Rep ΔC33 or K28A mutants, no removal of the roadblock or active replication is observed (Figures 3E, F, and G, Supplementary Figure S7). The experimental setup of these single-molecule reactions involves only pre-incubated DnaBC with the template DNA, and DnaBC is not present free in solution during the replication reaction. Therefore, these results suggest that the DnaB helicase remains bound after the replisome encounters the dCas9-cgRNA1-Atto647 roadblock, consistent with previous ensemble *in vivo* studies (64–66). Taken together, these observations demonstrate that Rep WT effectively removes the dCas9-cgRNA1 complex and that the success of this activity is dependent on a functional ATPase domain and the presence of the C-terminus that interacts with DnaB.

### Rolling-circle DNA templates show periodic replication stalling and rescue events

The rolling-circle DNA template allows for the observation of replication events that proceed for extended periods, limited only by the amount of nucleotides in the solution. Therefore, we hypothesized that the introduction of both the dCas9 roadblock and Rep in solution would result in the observation of multiple cycles of dCas9 binding, fork stalling and Rep-mediated rescue on individual molecules. Here, we repeated rolling-circle DNA replication assays with two modifications; (1) adding Rep-AF647 and dCas9-cgRNA1 complexes in solution with the replisome components, and (2) using an 18-kbp rolling-circle DNA template to resolve stalling and rescue events unambiguously (Figure 4A). The 18-kbp rolling-circle DNA template is capable of producing long DNA products at a rate similar to that of the 2-kbp rolling-circle DNA template (Supplementary Figure S8). We show in bulk biochemical assays, using the 2-kbp rolling-circle DNA template, that the Rep-AF647 protein is successful in removing the dCas9-cgRNA1 complex, an activity that is increased in the presence of increased ATP concentration under the conditions used (Supplementary Figure S9). The addition of Rep-AF647 and dCas9-cgRNA1 complexes to the single-molecule 18-kbp rolling-circle DNA replication assay resulted in long DNA products with multiple pausing and rescue events, where trajectories resembled steps at the expected binding sites of the roadblock (Figure 4B and Supplementary Figure S8B).

**Figure 4.**
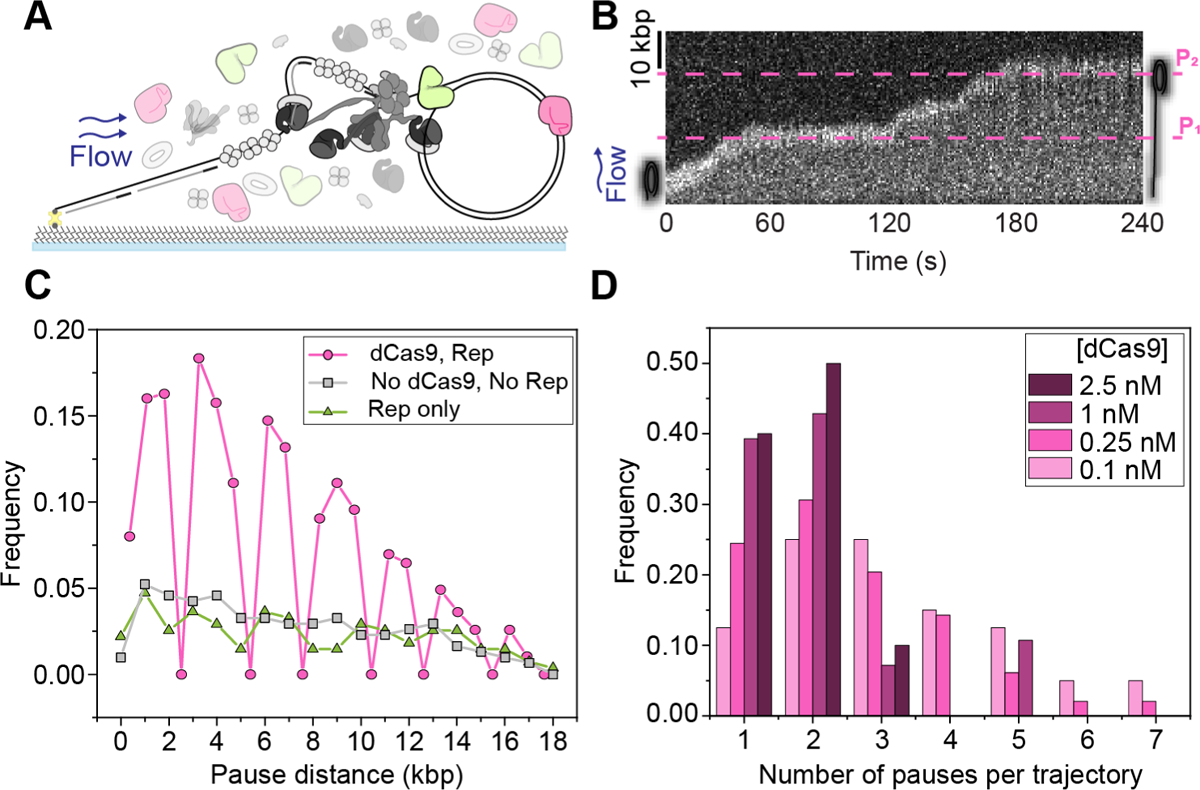
Observations of multiple stalling events. **(A)** Schematic representation of single-molecule stalled replication rescue assays, pre-incubated with DnaBC and immobilized to the flow cell surface. Replication is initiated in the presence of Rep and dCas9-cgRNA1. **(B)** Example 18-kbp rolling-circle DNA template undergoing multiple replication stalling and rescue events, at approximately 17 kbp (P_1_) and 36 kbp (P_2_). The target site of the dCas9-cgRNA1 complex occurs once every 18 kbp of the DNA template. **(C)** Pairwise distance analysis of the pause start sites replication rescue events on the 2-kbp DNA template in the presence (magenta) of 10 nM Rep-AF647 and 0.25 nM dCas9-cgRNA1 (16 pauses/275 kbp), only Rep-AF647 (green) (16 pauses/275 kbp) and absence of both proteins (gray) (18 pauses/307 kbp), for the first 20 kbp of DNA products. Symbols represent the distribution of histogram bin heights, normalized to the total DNA product length. Pauses in the absence of dCas9-cgRNA1 represent spontaneous pausing of the replisome. **(D)** Histograms of the number of pauses per replicating molecule at titrated dCas9-cgRNA1 complexes in the presence of 10 nM Rep-AF647 using the 2-kbp rolling-circle DNA template; 2.5 nM dCas9-cgRNA1, 17 pauses from 10 products (replication efficiency of 0.9 ± 0.3% (mean ± S.E.M.)); 1 nM dCas9-cgRNA1, 56 pauses from 30 products (1.4 ± 0.2 %); 0.25 nM dCas9-cgRNA1, 128 pauses from 51 products (3.0 ± 0.3%); and 0.1 nM dCas9-cgRNA1, 130 pauses from 43 products (2.7 ± 0.3%).

We performed change-point analysis (52, 53) of the trajectories of the position of the replicating DNA molecule in the movies using an automated tracking algorithm. Here, the individual rate segments were defined as stalled replication events or pauses, where the replication rate was below 100 bp s^-1^. Using this definition, we could then determine the pause sites (expressed in kbp from the start of the replication reaction). Using the 2-kbp rolling-circle DNA template, we observed multiple stalling and rescue events in reactions containing dCas9-cgRNA1 and Rep in solution (Supplementary Figure S10A). To confirm these observations, we repeated the experiment with the 18-kbp rolling-circle DNA template, which showed pausing at the expected target sites of the dCas9-cgRNA1 complex and subsequent rescue (Figure 4B and Supplementary Figure S8B). Pairwise distance analysis of the pause start sites of the 2-kbp rolling-circle DNA templates shows that replisome stalling and rescue events occur at every 2 kbp, or integers of 2 kbp, evident by the clustering around these distances and consistent with our expectations (Figure 4C).

In the absence of dCas9-cgRNA1 complexes, the periodic pausing behavior of the replisome was absent. Spontaneous pausing of DNA replication was observed to occur at any site, regardless of the presence of Rep-AF647. Periodic replication stalling and rescue was also observed in assays containing dCas9-cgRNA4 complexes targeted to the leading-strand of the 2-kbp rolling-circle DNA template (Supplementary Figure S10B). Despite the lower spatial precision in pause-site identification in the 18-kbp rolling circle template experiments, clusters in the periodicity of pausing are observed only in experiments containing both dCas9-cgRNA1 and Rep-AF647 (Supplementary Figure S8D). Having confirmed that the replication pausing occurs at the dCas9 binding sites, remaining experiments were conducted using the 2-kbp rolling-circle DNA template due to the low experimental throughput of the 18-kbp template.

We next titrated dCas9-cgRNA1 in solution to observe the effect of the number of stalling and rescuing events per replicating molecule. We observed a significant decrease in both the number of stalling and rescue events and the efficiency of molecules undergoing replication at high (1–2.5 nM) dCas9-cgRNA1 concentrations (Figure 4D). Therefore, subsequent reactions were conducted at lower dCas9-cgRNA1 concentrations.

### Resolution of stalled replication by Rep shows one rate-limiting step

Having established that we can observe multiple pause and rescue events using the 2-kbp rolling-circle DNA template, we next set out to investigate the activity of Rep during the pause states of the replication fork. We first identified pauses in rolling-circle DNA replication reactions containing only 0.25 nM dCas9-cgRNA1 in solution with the replisome components, or 0 nM dCas9-cgRNA1 and 0 nM Rep-AF647. Notably, in reactions containing only 0.25 nM dCas9-cgRNA1 the pause duration was recovered from a Gaussian distribution to be 135 ± 61 s, almost thirty-fold higher than spontaneous pauses identified in reactions containing 0 nM Rep-AF647 and 0 nM dCas9-cgRNA1 (5 ± 2 s), recovered from a single-exponential distribution (Figure 5A). This striking increase in pause duration confirms that pauses observed in reactions containing dCas9-cgRNA1 are caused by the roadblock complex. It is important to note that the duration of pauses in the dCas9-cgRNA1 only conditions reflects the period of time where the replisome was paused after initiation of replication until the end of the 4-minute acquisition. Therefore, the duration of a pause in the absence of Rep is an underestimate, given that previous estimates of the lifetime of the roadblock is on the tens of hours scale (35).

**Figure 5.**
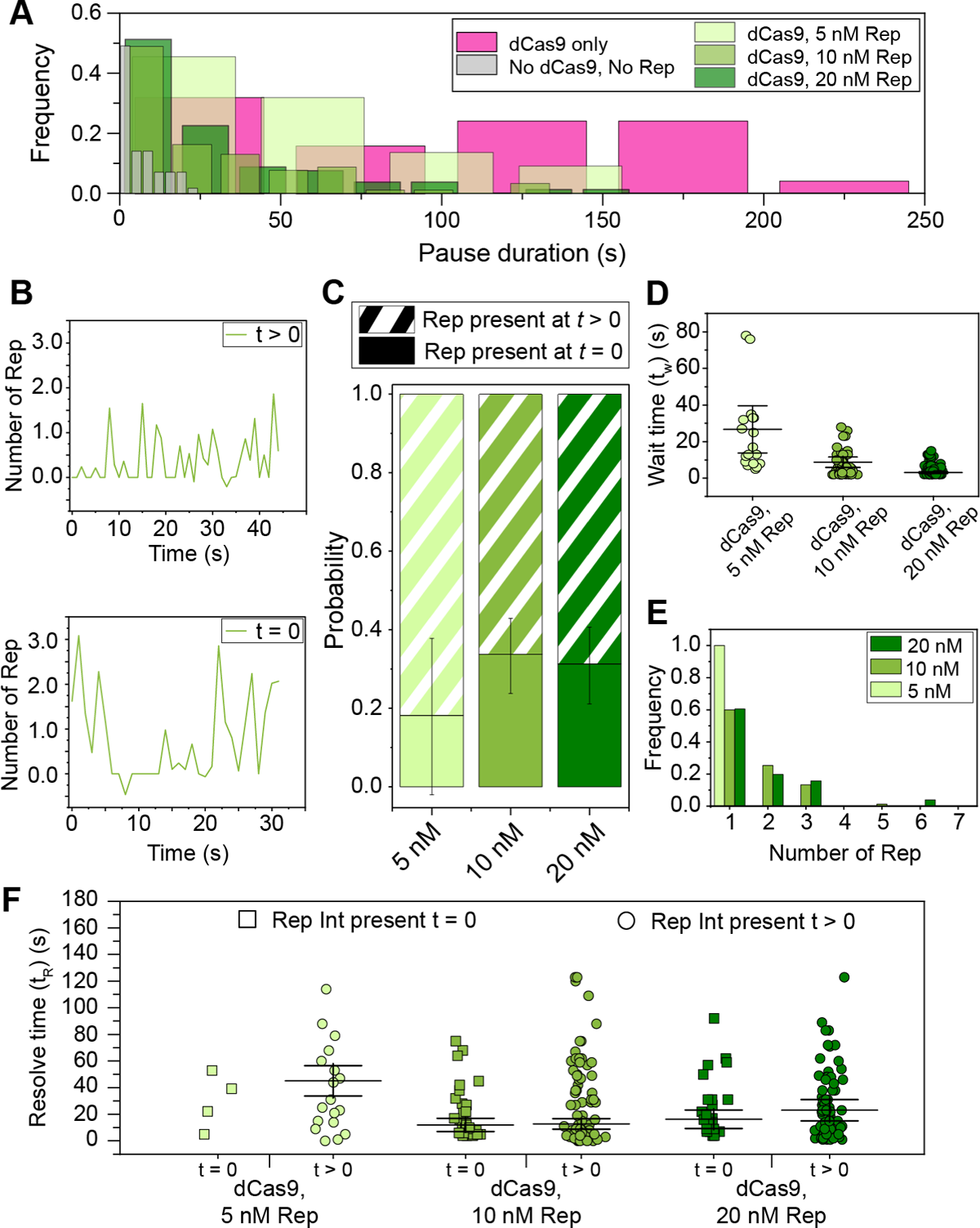
Observations of Rep at stalled replisomes. **(A)** Duration of a pause is decreased at increasing concentrations of Rep. In the presence of 0.25 nM dCas9 only, mean pause duration is determined from fitting a Gaussian distribution function (135 ± 61 s (S.E.M.), *n* = 25 pauses). In the presence and absence of dCas9-cgRNA1 and Rep-AF647, mean pause duration is determined by fitting single-exponential decay functions to the data (No dCas9-cgRNA1, No Rep-AF647, 5 ± 2 s, *n* = 57 pauses; dCas9-cgRNA1 and 5 nM Rep-AF647, 77 ± 38 s, *n* = 22 pauses; dCas9-cgRNA1 and 10 nM Rep-AF647, 21 ± 7 s, *n* = 92 pauses; dCas9-cgRNA1 and 20 nM Rep-AF647, 24 ± 9 s, *n* = 80 pauses). Pause durations of titrated Rep conditions represent pauses where a Rep-AF647 intensity was observed above a threshold. **(B)** Example traces of the number of Rep-AF647 present during a pause as a function of time. Two distinct types are observed; Rep-AF647 intensity above the threshold is reached at *t* > 0 (top), and Rep-AF647 intensity above the threshold is reached at *t* = 0. **(C)** The probability of observing the two Rep-AF647 activities during a pause for each concentration of Rep. Error bars indicate the margin of error for each concentration. **(D)** Distribution of the wait time (*t*_w_) for a Rep-AF647 molecule to associate to the replication fork in *t* > 0 events. Mean wait time for each concentration of Rep-AF647 was determined from fitting a single-exponential decay function to the data: 5 nM Rep-AF647, 27 ± 38 s (*n* = 18 pauses); 10 nM Rep-AF647, 8.7 ± 3.1 s (*n* = 76 pauses); 20 nM Rep-AF647, 2.7 ± 0.8 s (*n* = 78 pauses). **(E)** Histogram showing distributions of the number of Rep-AF647 molecules that associated first during a pause at *t*_w_, reveal predominantly monomeric stoichiometry at all concentrations used. **(F)** Pause resolve time (*t*_R_) for *t* = 0 (squares) and *t* > 0 (circles) events. The mean pause resolve times were determined by fitting a single-exponential decay function to the data of each concentration: 5 nM Rep-AF647, (*t* = 0) no fit converged (*n* = 4 pauses), and (*t* > 0) 46 ± 58 s (S.E.M.); 10 nM Rep-AF647, (*t* = 0) 11.7 ± 1.7 s (*n* = 31 pauses), and (*t* > 0) 12.5 ± 4.0 s; 20 nM Rep-AF647, (*t* = 0) 17 ± 7 s (*n* = 25 pauses), and (*t* > 0) 24 ± 39 s.

Next, we titrated Rep-AF647 in solution with dCas9-cgRNA1 complexes and visualized it at sites of stalled replication forks (Supplementary Figure S10A). In these dual channel videos, we identified pausing of the DNA replisome in the Sytox channel, and measured the lifetime of the pause where association of Rep-AF647 to the site was detected. We observed a reduction in pause duration with increasing concentrations of Rep-AF647. Specifically, at 5 nM Rep-AF647 the pause distribution was best described by a single-exponential fit with a mean duration of 77 ± 38 s, that reduced to 21 ± 7 s and 24 ± 9 s at 10 and 20 nM Rep-AF647, respectively. The plateauing of the pause duration at approximately 20 s for both the 10 and 20 nM Rep-AF647 conditions, suggests a saturating concentration has been reached. The significant reduction in pause duration when compared to the dCas9-cgRNA1 only conditions provides evidence that Rep-AF647 is required to remove the roadblock. Further, the single-exponential distributions of the pause durations suggest there exists a single rate-limiting step governing replication restart.

To understand the behavior of Rep-AF647 during these pause events, we plotted the intensity profiles of Rep-AF647 over time during the pause. This revealed two distinct types of Rep-AF647 signals: (1) events in which Rep associated with the replication fork after the pause started (*t* > 0) or (2) events in which Rep-AF647 was already present at the replication fork when the pause event began (*t* = 0) (Figure 5B). The likelihood of these two distinct events occurring was also dependent on the concentration of Rep-AF647 in solution: at higher Rep-AF647 concentration we detected a greater fraction of events in which Rep-AF647 was present at *t* = 0 (5 nM = 18 ± 20%; 10 nM = 34 ± 10%; 20 nM = 31 ± 10%) (Figure 5C).

Additionally, we quantified the average binding frequency of Rep-AF647 during the pause state of the replication fork to be slightly higher than that of an elongating replication event. At 20 nM Rep-AF647 a binding frequency of 22 ± 2 events per minute of stalled replication (mean ± S.E.M., *n* = 132 pauses) was determined. This frequency decreases at 10 and 5 nM Rep-AF647, where binding frequencies of 16 ± 1 (*n* = 128 pauses) and 2.0 ± 0.2 (*n* = 132 pauses) events per minute were determined, respectively. The reported binding frequencies during a pause are slightly higher than observed during elongation at the same concentrations (20 nM, 16 ± 1 min^-1^; 10 nM, 10 ± 1 min^-1^; and 5 nM, 7 ± 1 min^-1^) These observations rule out a scenario in which Rep is specifically recruited to the stalled replisome following an encounter with a roadblock; rather the association of Rep with the replication fork is stochastic and more frequent at higher concentrations of Rep in solution.

We further analyzed the events where Rep associates at *t* > 0. These events provide the opportunity to determine the characteristics of the associating Rep molecules after the replisome has stalled, rather than the Rep molecules that were already present at the fork when stalling occurred. We quantified the wait time (*t*_w_) for Rep-AF647 molecules to associate with the stalled replication fork using the increase in the intensity of Rep-AF647 molecules co-localizing with the fork at *t* > 0 (Supplementary Figure S10E). As expected, the *t*_w_ for associating Rep-AF647 molecules decreased with increasing Rep-AF647 concentrations, where single-exponential fit to the distribution of wait times revealed a mean *t*_w_ of 27 ± 38, 8.7 ± 3.1, and 2.7 ± 0.8 s for 5, 10, and 20 nM Rep-AF647, respectively (Figure 5D). These waiting times reflect the association rate constant of Rep to the replisome as the wait time decreases with increasing Rep concentrations.

Further, we quantified the stoichiometry of the first associating Rep-AF647 molecules during the pause states. Our assays revealed that the predominant stoichiometric state of an associating Rep-AF647 molecule is the monomer at all concentrations of Rep-AF647 used (Figure 5E). Our observations of two or more Reps binding could represent either the binding of higher oligomeric states, or the association of multiple monomers to the DnaB helicase. Under the experimental conditions used, the intensity quantification is robust enough to see differences between monomers and dimers, but not sufficient to distinguish between dimers or higher oligomers.

Finally, we quantified the pause resolution time (*t*_R_) for both types of Rep-association, where *t*_R_ is defined as the duration between the time at which the Rep-AF647 intensity exceeds the background and the time at which the pause is resolved in the corresponding DNA channel (Supplementary Figure S10E). This time period corresponds to the activities needed for roadblock removal and re-start of replication elongation. Interestingly for both observed categories, the pause resolution times were similar at each concentration, especially at 10 nM Rep-AF647 where *t*_R_ was determined from single-exponential fits to the resolution time distributions to be 12.5 ± 4.0 s and 11.7 ± 1.7 s for *t* > 0 and *t* = 0, respectively (Figure 5F). This similarity, in addition to the single-exponential distributions, provides further evidence that a single rate-limiting step governs the rescue of stalled replication upon Rep association. We cannot distinguish, however, whether this rate-limiting step corresponds to roadblock removal or another process underlying the re-start of replication. Taken together our results support a scenario in which Rep interrogates the state of the replication fork through frequent and stochastic association with the replisome.

### Pause duration does not depend on the stability of the roadblock

Having observed that Rep-AF647 can effectively remove dCas9-cgRNA1 complexes that are fully complementary to the DNA target, we set out to investigate the effect of roadblock stability on the pause duration. dCas9-cgRNA1 complexes bind their target sites tightly with 75% of complexes remaining bound after 16 h (35). The number of mismatched gRNA-target DNA bases has a significant effect on the stability of the complex (67–70). Specifically, multiple mismatches in the PAM-distal end of the gRNA-target DNA hybrid trigger faster dissociation. Therefore, we designed a set of mismatch (MM) cgRNAs containing 20–80% complementarity to the original target on the lagging strand of the 2-kbp rolling-circle template (termed cgRNA80, cgRNA60, cgRNA40, and cgRNA20) (Supplementary Figure S11A).

First, we estimated the approximate binding lifetime of these MM cgRNAs to an 83-mer target dsDNA sequence containing a single target site in single-molecule TIRF assays (Figure 6A). Here, the 83-mer target dsDNA was immobilized to the coverslip surface and videos of Sytox orange-stained dsDNA were collected as dCas9-MMgRNA-Atto647 was introduced to the flow cell. Images were acquired intermittently every 30 s for a total of 10 min. We observed efficient and stable binding of the dCas9-cgRNA80-Atto647 complex until the end of the acquisition, with a mean binding lifetime of 8.60 ± 0.03 min (mean ± S.E.M., *n* = 1414) (Figure 6B and Supplementary Figure S11B). This binding lifetime was similar in complexes containing 60% complementarity (cgRNA60, 8.30 ± 0.05 min, *n* = 619), where some complexes were observed to remain bound until the end of the acquisition, where others dissociated. Interestingly, complexes containing 40 and 20% complementarity showed very weak binding and were often only observed bound to the 83-mer target DNA for one frame (cgRNA40, 1.00 ± 0.05 min, *n* = 249; cgRNA20, 1.00 ± 0.06 min, *n* = 109). This is consistent with observations of dCas9 stability where cgRNA-target DNA hybrids containing less than 50% complementarity, especially in the reversibility-determining region, showed significantly reduced lifetimes (67) (Supplementary Figure S11A).

**Figure 6.**
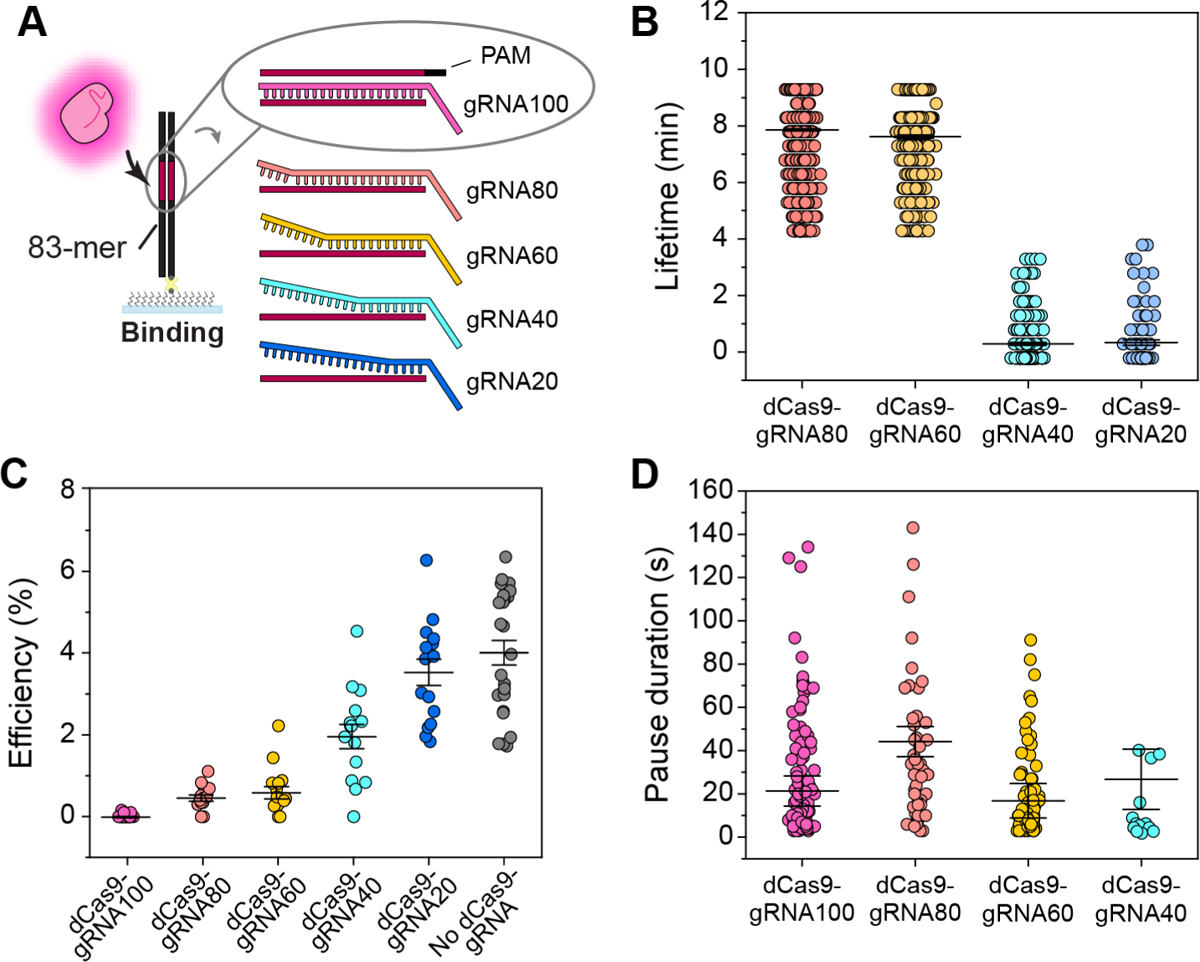
Stalled replication rescue at less stable roadblocks. **(A)** Schematic representation of dCas9-cgRNA mismatch complexes binding to 83-mer dsDNA oligonucleotides in single-molecule lifetime assays. **(B)** Lifetime distributions of dCas9-MMgRNA complexes binding to 83-mer dsDNA. Mean lifetime for each complex; dCas9-gRNA80, 8.60 ± 0.03 min (S.E.M.) (*n* = 1414 events); dCas9-gRNA60, 8.30 ± 0.05 min (*n* = 619 events); dCas9-gRNA40, 1.00 ± 0.05 min (*n* = 249 events); dCas9-gRNA20, 1.00 ± 0.06 min (*n* = 103 events). **(C)** The efficiency of replication distributions of DNA templates following pre-incubation with dCas9-gRNA complexes. Mean efficiencies for each complex; dCas9-gRNA100, 0.02 ± 0.01%; dCas9-gRNA80, 0.50 ± 0.07%; dCas9-gRNA60, 0.60 ± 0.1%; dCas9-gRNA40, 2.0 ± 0.3%; dCas9-gRNA20, 3.5 ± 0.3%; No dCas9 complexes, 4.0 ± 0.3%. **(D)** Pause duration distributions in the presence of dCas9-MMgRNA complexes and Rep-AF647. Mean pause durations were determined from fitting a single-exponential decay function to the data of each complex: dCas9-gRNA80, (44 ± 16 s (S.E.M.), *n* = 51 pauses, replication efficiency of 3.2 ± 0.3%); dCas9-gRNA60, (17 ± 6 s, *n* = 72, 3.3 ± 0.2%); dCas9-gRNA40, (28 ± 14 s, *n* = 16, 4.3 ± 0.6%). dCas9-gRNA100 data are duplicated from Figure 5A; dCas9-cgRNA1 and 10 nM Rep-AF647. Pause durations represent pauses where a Rep-AF647 intensity was observed above the threshold.

Next, we measured the efficiency of DNA replication on templates that were pre-incubated with the various dCas9-MMgRNA complexes. The 2-kbp rolling-circle DNA templates in the absence of roadblocks show a mean efficiency of 4% under the conditions used (Figure 6C). This is significantly reduced when DNA templates are pre-incubated with dCas9-cgRNA1 complexes (0.02%). Interestingly, dCas9-MMgRNA complexes containing 80 or 60% complementarity to the target exhibited a 2-fold increased efficiency of replicating products, in comparison to fully complementary complexes (cgRNA80, 0.5 ± 0.1%; cgRNA60, 0.6 ± 0.1 %). The efficiency was further increased in experiments containing dCas9-MMgRNA complexes with 40 and 20% complementarity (cgRNA40, 2.0 ± 0.3%; cgRNA20, 3.5 ± 0.3%). This is consistent with bulk biochemical assays where DNA replication products were only detected in reactions containing dCas9-MMgRNAs of 60-20% complementarity, and the absence of the block band at approximately 2.5 kbp, at 40 and 20% complementarity (Supplementary Figure S11C).

Finally, we investigated the duration of pauses resolved by Rep-AF647 when caused by dCas9-MMgRNA complexes. Since we saw a reduced lifetime and similar efficiency to that of replication for cgRNA20, we used the 80–40% MMgRNAs in these assays. As before, Rep-AF647 and the specified dCas9-MMgRNA were added to the flow cell containing DnaBC-DNA to initiate the replication reaction. Pauses detected and resolved by Rep-AF647 revealed similar durations despite decreasing complementarity to the target DNA (Figure 6D). Specifically, the pause duration corresponding to all of MMgRNAs were similar to that of the dCas9-cgRNA1 complex (cgRNA80, 44 ± 7 s; cgRNA60, 17 ± 8 s; cgRNA40, 28 ± 14 s,). The similar pause durations of each of the cgRNAs in the presence of RepAF647 suggests that the removal of the roadblock is not the rate-limiting process during the rescue of stalled replication. Rather, these results suggest that the activity of Rep at the stalled replication fork is quick, and that a process involved with subsequent replication restart is the rate-limiting step.

## DISCUSSION

In this study, we set out to visualize the *E. coli* Rep helicase as it interacts with elongating and stalled replisomes. Our single-molecule observations of Rep binding to DNA confirm early investigations whereby the affinity of Rep to ssDNA is significantly decreased by ATP hydrolysis. We observed frequent and stochastic association of Rep in a predominantly monomeric state to the elongating replisome as it replicates DNA. Our investigations of Rep at the stalled replisome revealed that Rep removes dCas9-cgRNA roadblocks resulting in the rescue of stalled replication. Further, we showed that the resolution of replication stalled at high-stability roadblocks occurs with kinetics that can be described with a single rate-limiting step, regardless of whether Rep was already present at the fork at the onset of the stall or whether Rep associated after the stall. Finally, we show that the duration a replisome is stalled is constant at less stable roadblocks, indicating that the rate-limiting step is a process involved in the restart of replication, and not roadblock removal. Together, these results provide insight into the activity of Rep at the replisome and allow us to propose a model describing how Rep protects the replisome and acts in the context of roadblocks.

The main aim of this study was to observe the Rep helicase at elongating and stalled replisomes. While Rep is known to interact with the replisome, the context of this interaction is not well defined. Specifically, is Rep only present at the replisome in the stalled state, or is Rep continually interacting with the replisome? We show here that Rep interacts stochastically with the elongating replisome in the absence of protein roadblocks. The addition of Rep-AF647 into single-molecule rolling-circle replication assays shows that Rep can interact with the replisome and have no effect on its rate or processivity (Figure 2 and Supplementary Figure S4). Recent live-cell fluorescence studies proposed that Rep interacts with the replisome in low copy numbers or is only recruited to DnaB upon fork arrest (36). Here, we show that Rep frequently interacts with the elongating replisome in a predominantly monomeric stoichiometry at all concentrations used (Figure 2D). However, the frequency of binding to the replisome increases with increasing concentrations of Rep, suggesting this interaction is stochastic.

The stoichiometry of Rep at the replisome has been hypothesized to be hexameric, assuming all sites are occupied on the hexameric DnaB (36, 58). Our studies, both in the absence and presence of roadblocks, reveal a monomeric stoichiometry when Rep is associated with the replisome (Figure 2D and Figure 5E). Recent single-molecule live-cell studies observed up to six Rep monomers at the replication fork (24). At the concentrations used in our assays, we are well below the predicted micromolar cellular concentration of Rep (24). However, previous studies have also predicted that Rep may be present at the replisome in less than 3 copies (36). Together, these results show the plasticity of the replisome and could suggest that while Rep could occupy all binding sites on DnaB, the likelihood of this occurring could be dependent on other DnaB interacting partners. Previous surface plasmon resonance investigations estimated the apparent *K*_D_ of the Rep-DnaB interaction to be approximately 90 nM (17). However, these results are obtained outside the context of the functional replisome. Without structural information on the Rep-DnaB interaction, it cannot be certain that Rep does not interact in the same binding pockets as other essential components of the replisome. Therefore, we predict that the association of Rep to DnaB, and thus our reported stoichiometry, occurs not only due to the low concentrations used but also due to shared binding sites becoming available during replication.

Rep removes protein roadblocks from the path of the replication fork. In our study we observe the robust displacement of the dCas9-cgRNA roadblock in ensemble and single-molecule assays, resulting in the restart of replication. Pre-incubation of the rolling-circle DNA template with the dCas9-cgRNA1 roadblock showed a clear restart of replication after the disappearance of the roadblock fluorescent signal (Figure 3). This activity was not observed in the presence of either the ATPase deficient Rep K28A or DnaB interaction deficient Rep ΔC33 mutants, thus indicating that both activities are required for the removal of protein roadblocks from the template DNA. This observation is in agreement with previous studies that showed that Rep mutants lacking these structural components could not displace RNAP or other model roadblocks (17). Additionally, single-molecule live-cell studies showed that the C-terminus is required for the association to the replication fork, while the ATPase activity is required for translocation away from the fork (24). Recent studies have also shown that the 2B subdomain of Rep is essential for protein roadblock displacement (32, 33). Therefore, it is likely that both the interaction with the DnaB helicase, the functional ATPase domain, and the 2B subdomain are all essential to the displacement of roadblocks by Rep.

We report here the first real-time observations of protein displacement and the rescue of stalled replication by Rep. The single-molecule rolling-circle replication assays containing both Rep-AF647 and dCas9-cgRNA1 roadblocks in solution revealed a pause duration dependent on the concentration of Rep-AF647 (Figure 5A). This observation suggests that the higher the local concentration of Rep, the quicker the resolution of the roadblock due to a shorter search time. Interestingly, we observed a plateauing of the pause duration at 10 and 20 nM Rep-AF647. Despite the predicted *K*_D_ of Rep-DnaB being much higher, this saturation suggests that the *K*_D_ of the Rep-DnaB interaction within a functional replisome might be significantly lower, potentially due to the availability of higher affinity binding sites upon replisome stalling. The predicted micromolar cellular concentration of Rep would suggest that pauses in cells are resolved quicker than in our reconstituted replisome experiments. Nonetheless, our assays provide insight into the mechanisms required for Rep to displace protein roadblocks and rescue stalled replication.

We observed two distinct classes of Rep-mediated roadblock removal events; (1) where Rep is already present at the replisome upon stalling, and (2) where Rep associates after the replisome stalls (Figure 5B, C). While the latter activity could suggest recruitment to the stalled replisome, both activities resulted in similar pause resolution times, suggesting that once Rep is present the displacement of the roadblock occurs at the same rate through the same process. Further, we observed that the wait time for Rep to associate with stalled replisomes was less at higher concentrations of Rep (Figure 5D). Unlike the observed pause durations, the reported wait times did not plateau. Therefore, we hypothesize that the Rep association to the stalled replisome is independent of the affinity of the binding site and represents the concentration-dependent bimolecular association rate. Further, we predict that at more biologically relevant concentrations, Rep would interact with the replisome more frequently, further decreasing potential association wait times. The observed single-exponential distributions throughout the pause duration and pause resolution time all provide evidence that there is a single rate-limiting kinetic step governing the rescue of stalled replication once Rep associates with the replisome.

Our investigation of Rep at the sites of roadblocks with decreased stability provided further evidence that there is one rate-limiting step of stalled replication rescue. Investigations of mismatched RNA-DNA hybrids in complex with dCas9 have shown that only 8 bp of complementarity is required to establish a stable complex (67). Interestingly, our investigations revealed comparable pause durations when the replisome was stalled by dCas9-MMgRNA complexes to fully complementary roadblocks despite the significantly lower extent of complementarity in the R-loop (Figure 6D). The intrinsic lifetimes of the MMgRNAs (Figure 6B) and the constant pause durations suggest that a slower process after the removal of the protein roadblock is the rate-liming step of the reaction. Further, these results suggest that the roadblock removal activity of Rep once associated with the stalled replisome, is relatively quick, occurring on the time scale of a few seconds.

Additionally, we detected higher replication efficiencies when DNA templates were pre-incubated with the less stable dCas9 complexes, in the absence of Rep (Figure 6C). While this is reflective of the observed lifetimes of these complexes, the higher efficiencies may also suggest that the replisome can bypass the less stable complexes without the need for Rep. Further investigations of the replisome, with or without Rep, at sites of stalled replication caused by unstable roadblocks will elucidate mechanisms of roadblock removal and bypass.

How does the frequent association of Rep to the replication fork result in roadblock removal? Our study allows us to propose a model of Rep activity at elongating and stalled replisomes (Figure 7). Our investigation provides evidence supporting a model proposed by a previous live-cell study, whereby Rep is associated with the replication fork during elongation (24). Further, we propose that the association of Rep to the replisome is stochastic and does not occur by a recruitment mechanism. Association of Rep through the interaction with the DnaB helicase allows Rep to frequently monitor the state of the replication fork, acting as a shield to potential roadblocks (Figure 7A). If a roadblock is detected or encountered, then displacement activity will result (Figure 7B). This activity is quick, whereas the processes required to restart replication are relatively slow (Figure 7C). Given that the rescue of stalled replication in our assays did not require helicase reloading mechanisms, we hypothesize that the time required to restart replication determines if DnaB will remain stable or if the entire replication fork will collapse. Previous ensemble *in vivo* studies of replication fork stalling have estimated that DnaB remains stable for up to 30 min after stalling (64–66).

**Figure 7:**
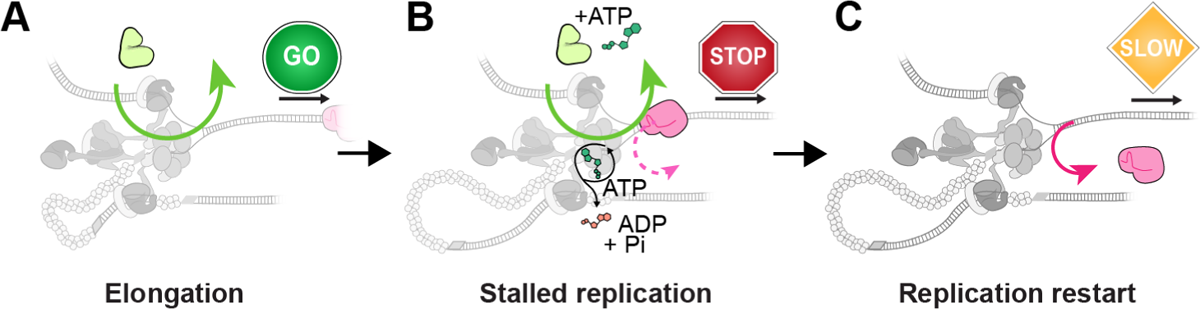
Model for Rep activity at elongating and stalled replication forks. **(A)** Rep stochastically associates with the replication fork during elongation. **(B)** Upon the replisome stalling, associated Rep molecules work to remove the roadblock from the path of the replication fork. This process is relatively quick. **(C)** Restart of replication after the removal of the roadblock is slow, representing the rate-limiting step in resolution of the stalled state.

Finally, our investigations also provide insight into the stability of the DnaB helicase when the replisome encounters a protein roadblock. Each single-molecule assay described in this investigation has DnaB pre-incubated with the DNA template and omits free DnaB complexes in solution during the replication reactions. The successful restart of replication after Rep has displaced the dCas9-cgRNA roadblock, without the need for additional protein factors, suggests that the pre-incubated DnaB helicase remains bound to the template DNA after an encounter with the roadblock. The DnaB helicase has recently been visualized in similar single-molecule assays to be a stable component of a processive replisome (39). While further investigations of the stability of other individual components of the replisome (for example, components of the Pol III holoenzyme and DnaG) are required, these recent observations suggest that the DnaB helicase is integral for replication restart. A stable DnaB helicase may act as a hub for the elongating replisome, allowing for efficient reloading of replisome components if collisions result and components of the replisome dissociate.

Single-molecule observations have proven valuable in elucidating the individual behaviors of replisome components (39,42,46). Our work, combined with other recent investigations, suggests a model where cooperation between Rep and the replisome is needed for efficient roadblock removal. We propose that Rep, and its homologs, fulfill their protective role for the replisome in an entirely stochastic manner that is not modulated by the replisome being in a stalled state. Further elucidation of both the Rep-replisome and the Rep-roadblock interactions will provide insight into the significant importance of this accessory helicase to cells.

## DATA AVAILABILITY

All data presented (DOI: 10.5281/zenodo.7375212) and home-built ImageJ plugins (DOI: 10.5281/zenodo.7379064) are available at Zenodo.org.

## ACKNOWLEDGEMENTS

We thank Prof. Timothy Lohman (WashU) for generously sharing expression vectors for Rep (A97C) used in these studies. We acknowledge the preliminary work by Finnian Fowke, Hamish Maynard, and Zhong Yan Gan. The authors thank Drs Gurleen Kaur, Richard Spinks and Stefan Mueller, and the wider van Oijen and Dixon groups for helpful discussions.

## Author contributions

Conceptualization: K.S.W., N.E.D., H.G., A.M.v.O.; methodology: K.S.W., Z-Q.X, S.J., N.S., H.G.; software: K.S.W., L.M.S.; validation: K.S.W., Z-Q.X., S.J.; formal analysis: K.S.W., S.J.; investigation: K.S.W., Z-Q.X., S.J., N.S.; resources: K.S.W., Z-Q.X., S.J. N.S.: writing: K.S.W., H.G., A.M.v.O.; visualization: K.S.W.; supervision: L.M.S., N.E.D., A.M.v.O., H.G.; funding acquisition: L.M.S., N.E.D., A.M.v.O., H.G.

## SUPPLEMENTARY DATA

**Supplemental Table S1:**
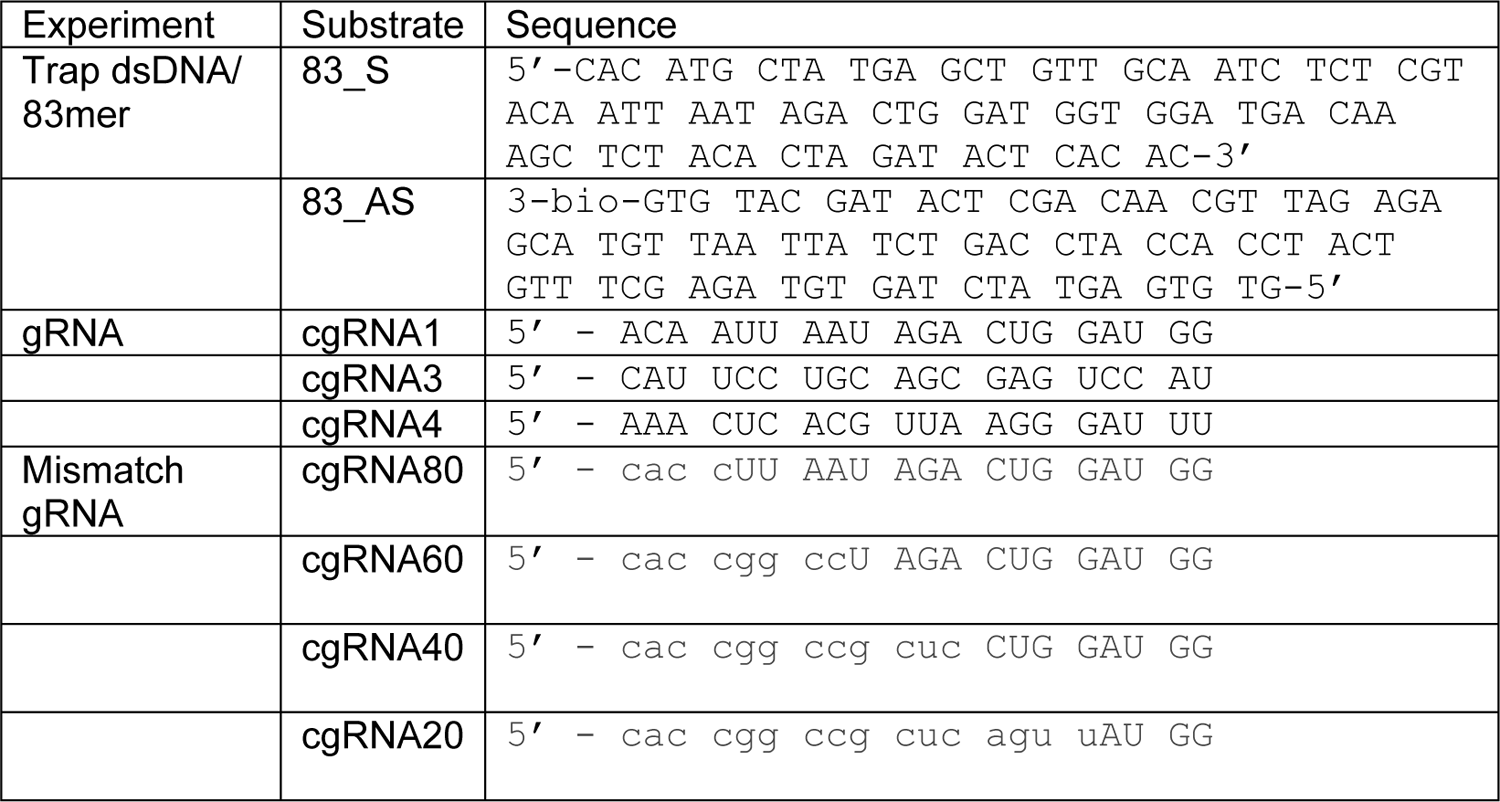
Nucleic acid substrates used in this study.

---

**Supplementary Figure S1:**
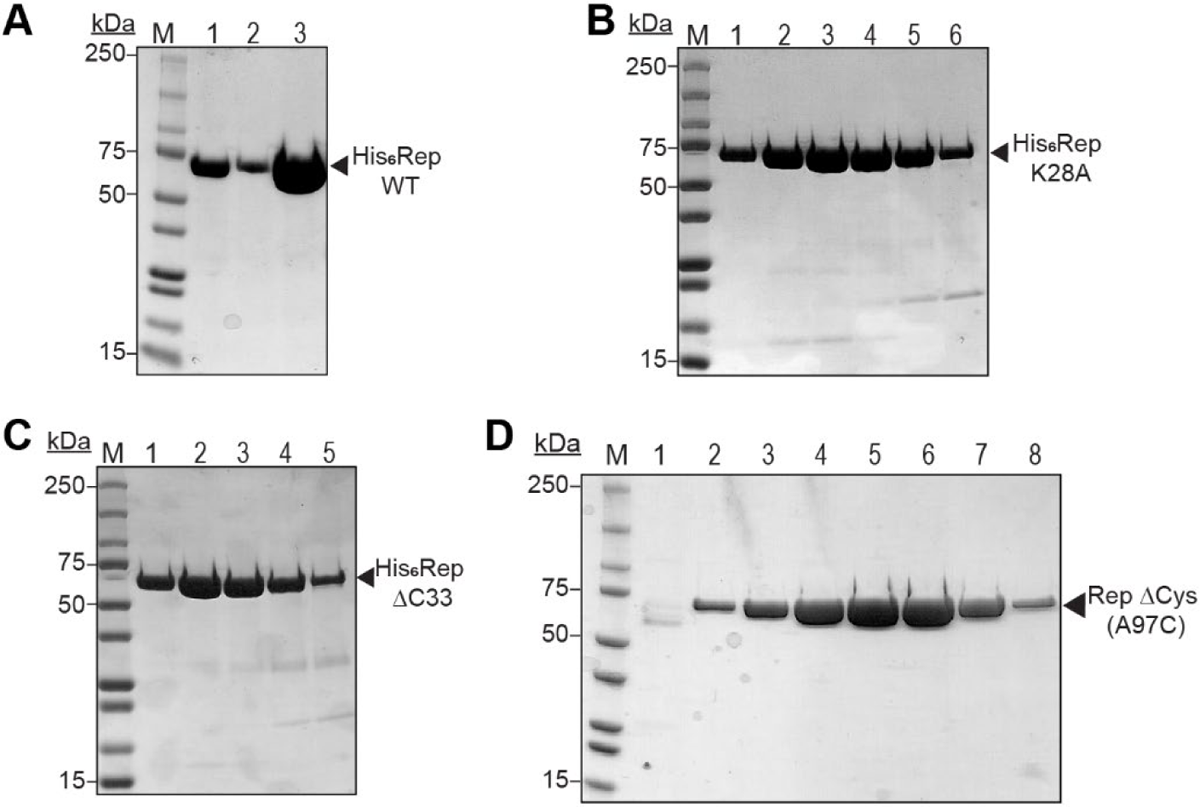
Purification of Rep proteins using 5 mL HiTrap Heparin columns. Samples were analyzed on 4–20% SDS-PAGE gels. **(A)** Purification of His_6_ Rep WT. Sample after purification on His-Trap column (lane 1), fractions of purified protein from Heparin column (lane 2–3) used in this study. **(B)** Purification of His_6_ Rep K28A. Samples from successive fractions (indicated by lane numbers 1–6). **(C)** Purification of His_6_ Rep ΔC33. Samples from successive fractions (indicated by lane numbers 1–5). **(D)** Purification of His_6_ Rep A97C. Samples from column flow-through (lane 1), and successive fractions (indicated by lane numbers 2–8).

**Supplementary Figure S2:**
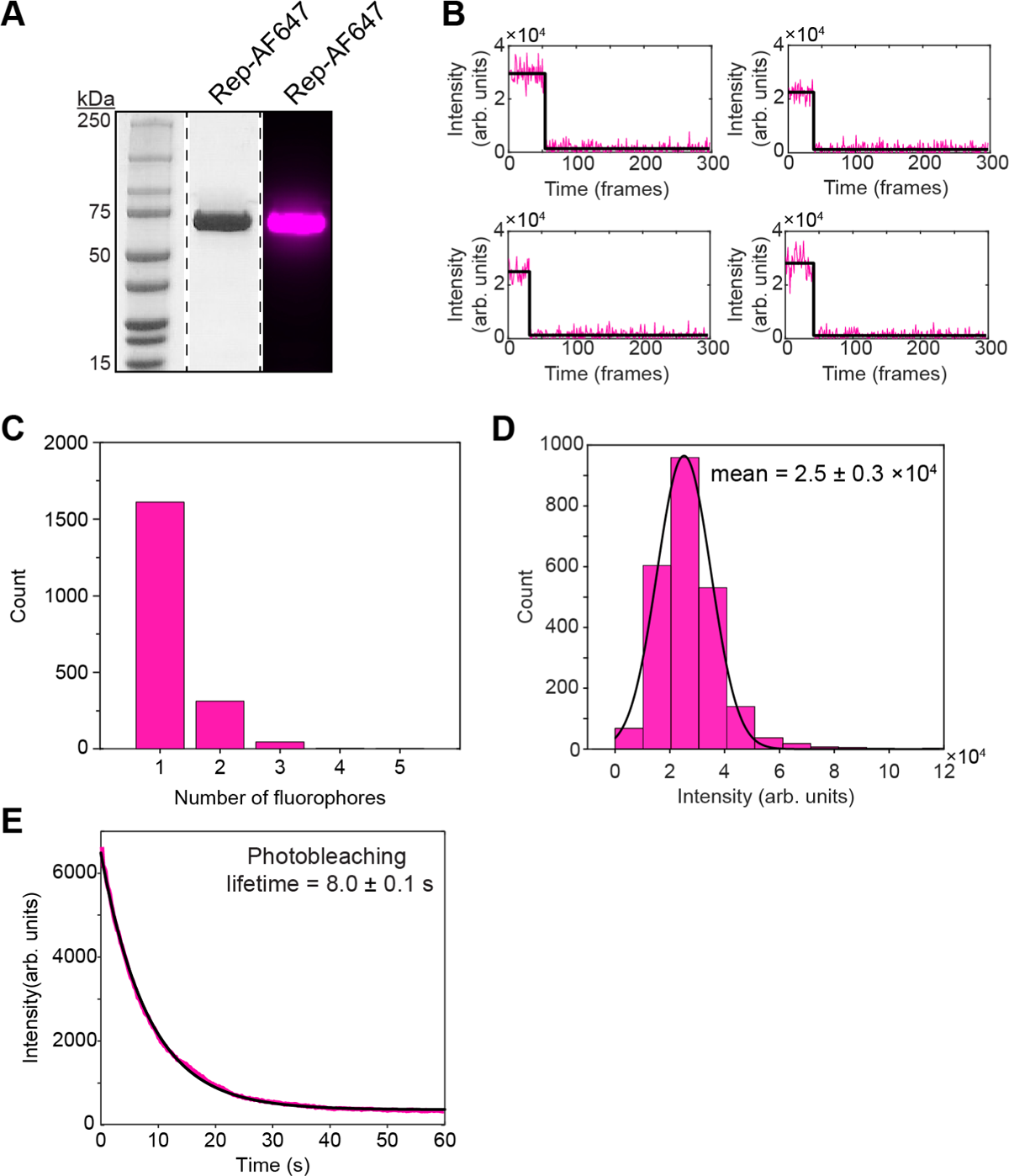
Quantification of fluorescent labeling of Rep-AF647. **(A)** SDS-PAGE gel of purified Rep-AF647. Left and middle lanes are stained with Coomassie blue and imaged using a Bio-Rad Gel Doc XR. Right lane is unstained and Alexa Fluor 647 fluorescence imaged using an Amersham Imager 600. **(B)** Example trajectories of Rep proteins deposited on coverslip and subjected to photo-bleaching. Number of fluorophores per Rep monomer are detected by quantifying the single-molecule photobleaching steps using change point analysis (black line). **(C)** Distribution of the number of steps **(B)** and therefore the number of fluorophores per monomer (*n* = 1483). **(D)** Distribution of the step size of (B) was 2.5 ± 0.3 x 10^4^ (mean ± S.E.M., *n* = 1483). **(E)** Average photo-bleaching trajectory for Rep-AF647 (*n* = 1483) at excitation power density of 200 mW cm^-2^. From a fit with a single-exponential decay function (black line), a photo-bleaching lifetime of 8.0 ± 0.1 s was obtained.

**Supplementary Figure S3:**
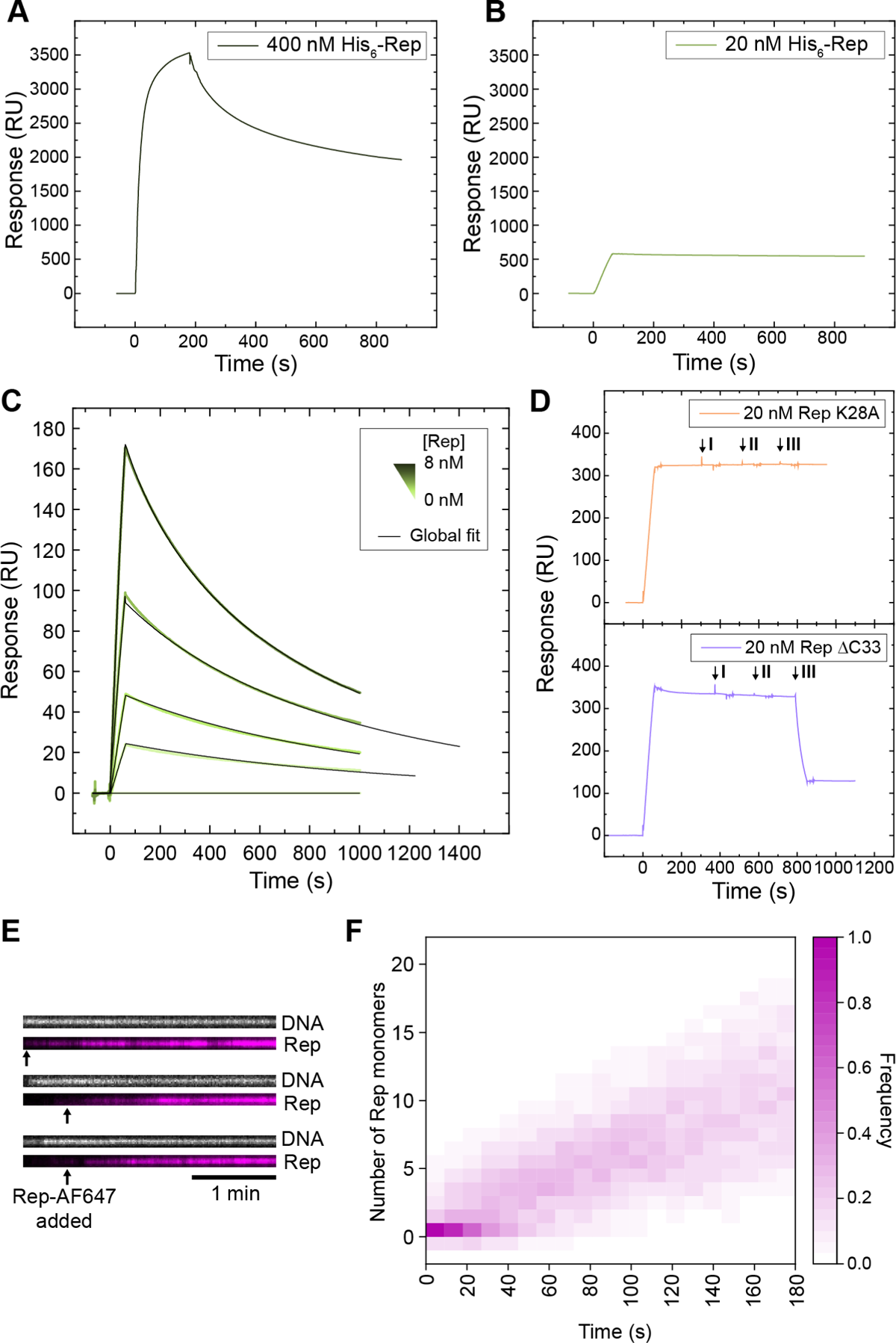
Observations of Rep binding to DNA. SPR sensorgrams of 400 nM. **(A)** and 20 nM **(B)** Rep WT binding to dT_35_. **(C)** Association (60 s) and dissociation of titrated (1–8 nM) Rep WT binding to dT_15_ (global fitting of Rep binding (1:1 binding with mass transfer) yielded a *K*_D_ of approximately 500 pM). **(D)** 20 nM Rep K28A (purple – top) and 20 nM Rep ΔC33 (orange – bottom) dissociation after nucleotide AMP-PNP (I), ADP (II) or ATP (III)) injection. **(E)** Example kymographs of Rep-AF647 binding to 2-kbp rolling-circle DNA template bound by SSB and DnaBC, in the absence of ATP. Arrows indicate the time point of addition of Rep-AF647 to the flow cell. **(F)** Heatmap of number of Rep-AF647 monomers bound to the DNA template over time in the absence of ATP (*n* = 70).

**Supplementary Figure S4:**
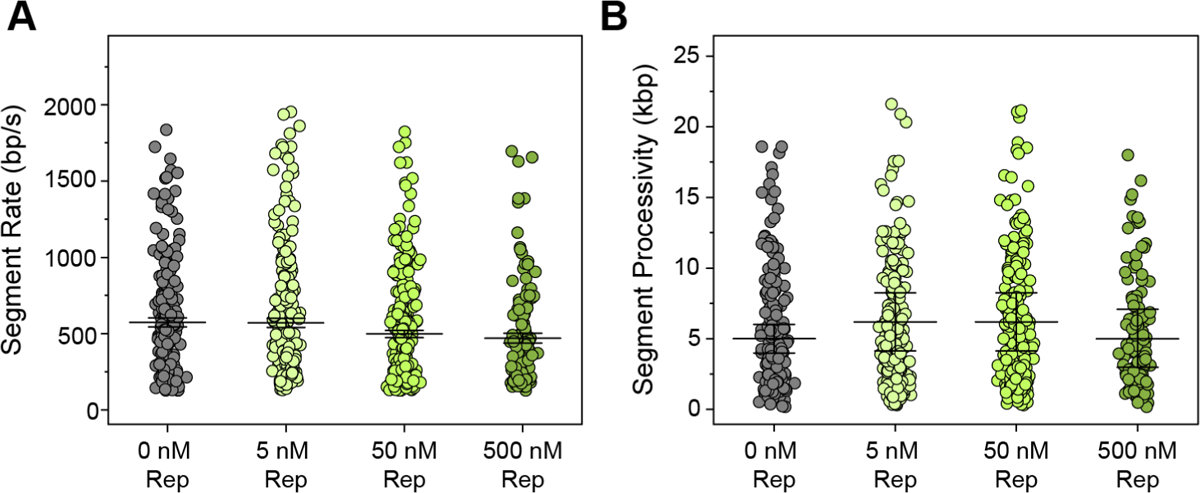
Quantification of the effect of Rep WT on the rate and processivity of replication. **(A)** Median rates of replication in the absence (gray) (576 ± 30 bp s^-1^ (median ± S.E.M., *n* = 179, replication efficiency = 5.2 ± 0.7% (S.E.M.)) and presence of titrated Rep WT (5 nM (light green) (573 ± 29 bp s^-1^ (*n* = 208, 4.5 ± 0.5%), 50 nM (olive green) (498 ± 24 bp s^-1^ (*n* = 234, 5.8 ± 0.7%), and 500 nM (dark green) (477 ± 32 bp s^-1^ (*n* = 113, 2.8 ± 0.4%)),quantified by change-point analysis of single-molecule rolling-circle DNA replication trajectories. **(B)** Mean processivity of replication in the absence (grey) (5 ± 1 kbp (mean ± S.D.)) and presence of titrated Rep WT (5 nM (light green) (6 ± 2 kbp), 50 nM (olive green) (6 ± 2 kbp), and 500 nM (dark green) (5 ± 2 kbp)). Mean processivities were determined from fitting a single-exponential decay function to data.

**Supplementary Figure S5:**
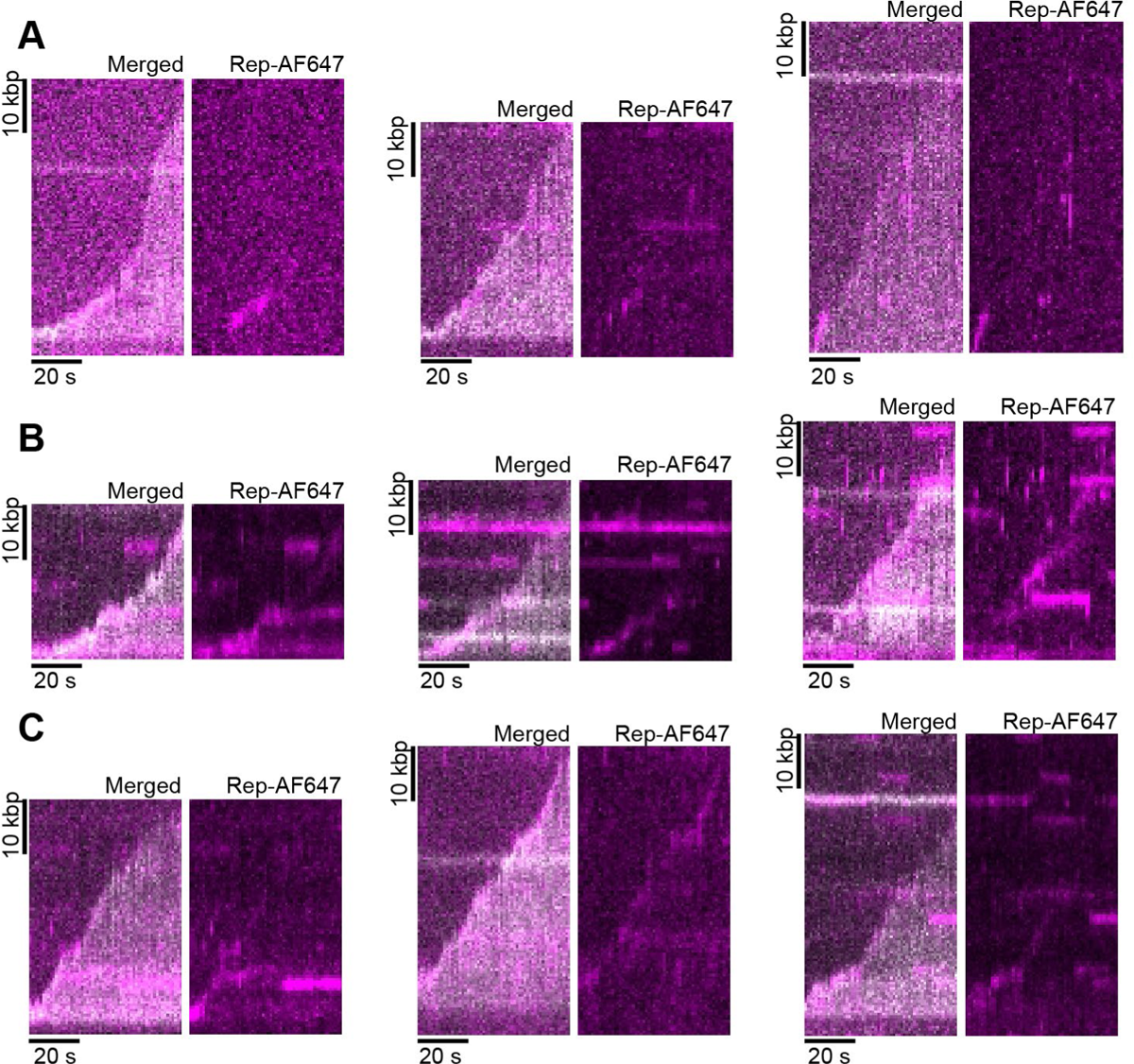
Example kymographs of Rep-AF647 during processive rolling-circle DNA replication. Merged kymographs of Rep-AF647 intensity (magenta) and Sytox orange-stained DNA (gray) (left) and Rep-AF647 intensity alone (right). **(A)** 5 nM Rep-AF647 (replication efficiency of 6.9 ± 0.3% (S.E.M.)). **(B)** 10 nM Rep-AF647 (5.4 ± 0.2%). **(C)** 20 nM Rep-AF647 (4.2 ± 0.2%).

**Supplementary Figure S6:**
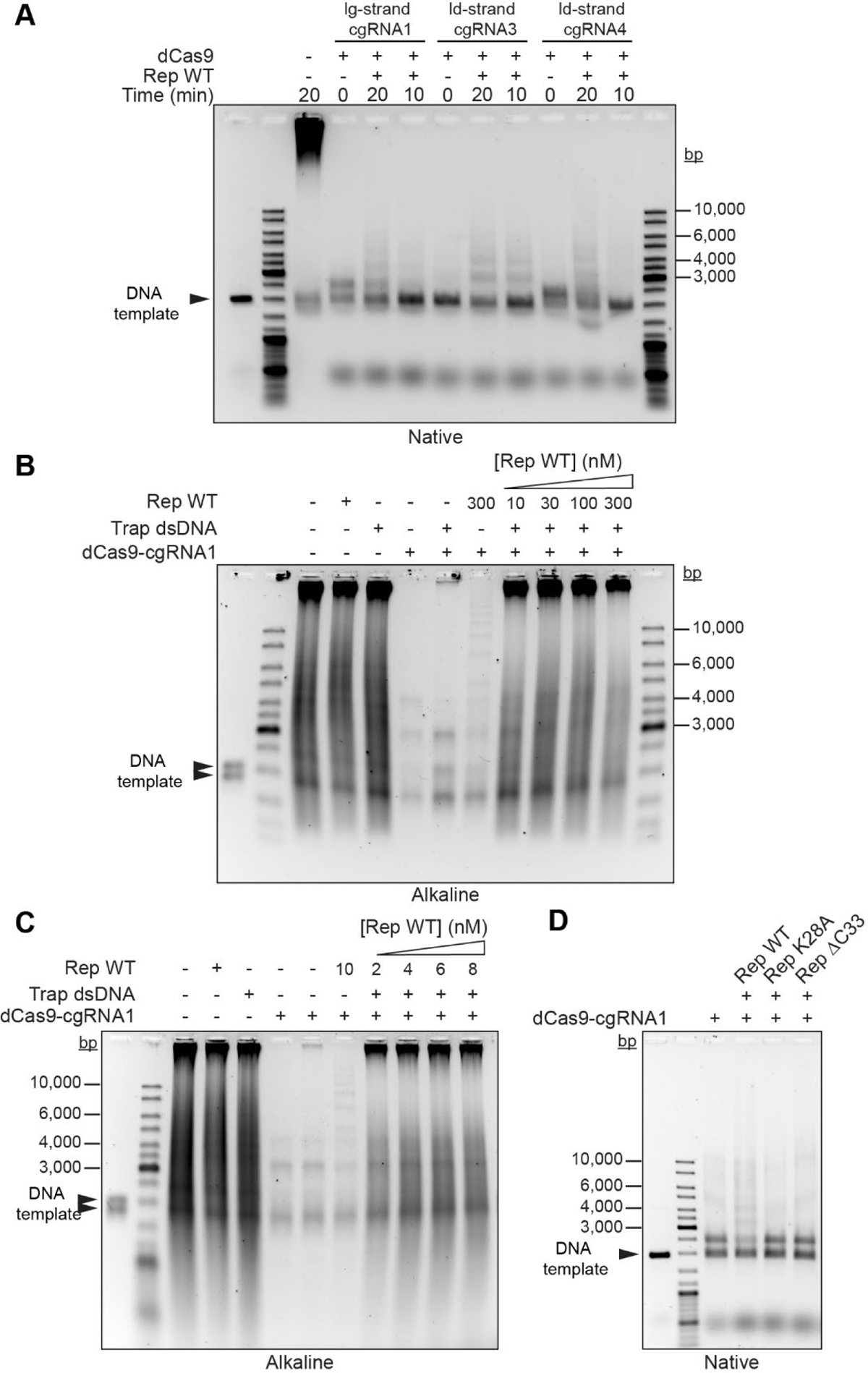
Rep rescues DNA replication stalled by dCas9-cgRNA complexes. Reactions contained 200 nM of specified cgRNA and 50 nM dCas9 complex. **(A)** Rep WT rescues stalled DNA replication independent of the DNA strand targeted by dCas9-cgRNA complex. Rep WT added to reactions at 10 min time point. *N* > 2 independent experiments. **(B)** Titration of Rep WT (10– 300 nM). Addition of trap dsDNA results in higher extent of rescued DNA products. *N* > 2 independent experiments. **(C)** Titration of Rep WT (2–8 nM). *N* > 2 independent experiments. **(D)** Rep mutants lacking either the C-terminal domain (ΔC33) or functional ATPase (K28A) cannot rescue dCas9-gRNA1 stalled DNA replication *N* > 2 independent experiments.

**Supplementary Figure S7:**
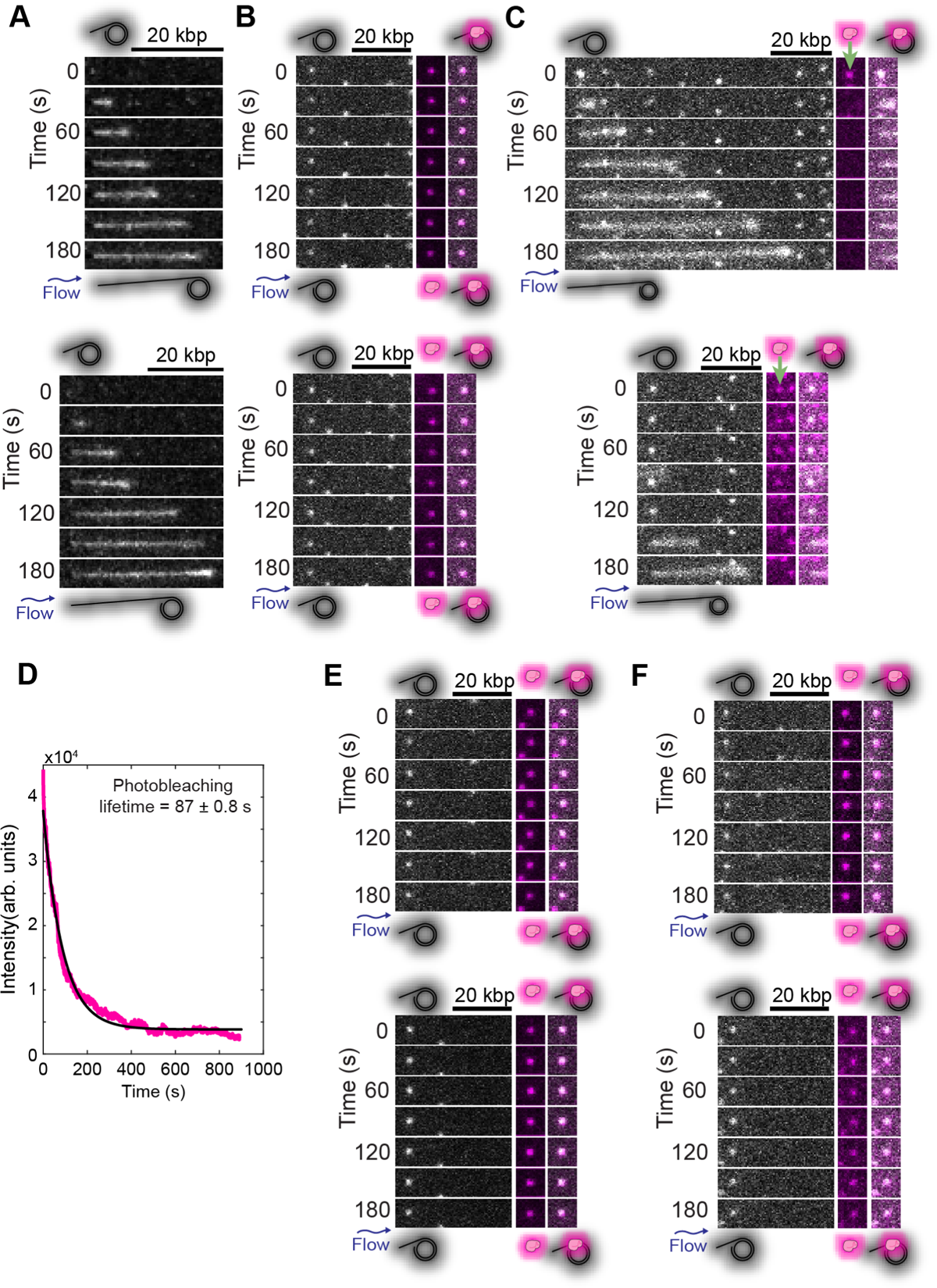
Additional example montages of replication rescue of DNA templates (Sytox orange stained; gray scale) pre-incubated with dCas9-cgRNA1-Atto647 (magenta) complexes. **(A)** In the absence of dCas9-cgRNA1 complexes. **(B)** Pre-incubation of the DNA template with dCas9-cgRNA1-Atto647. **(C)** Addition of Rep WT (20 nM) results in disappearance of dCas9-cgRNA1-Atto647 (green arrow). **(D)** Average photo-bleaching trajectory for dCas9-cgRNA1-Atto647 (*n* = 139) at excitation power density of 200 mW cm^-2^ imaged every 200 ms. Single-exponential decay fit (black line) revealed a photo-bleaching lifetime of 87 ± 1 s. **(E)** Rep K28A and **(F)** Rep ΔC33 cannot remove dCas9-cgRNA1-Atto647 complexes.

**Supplementary Figure S8:**
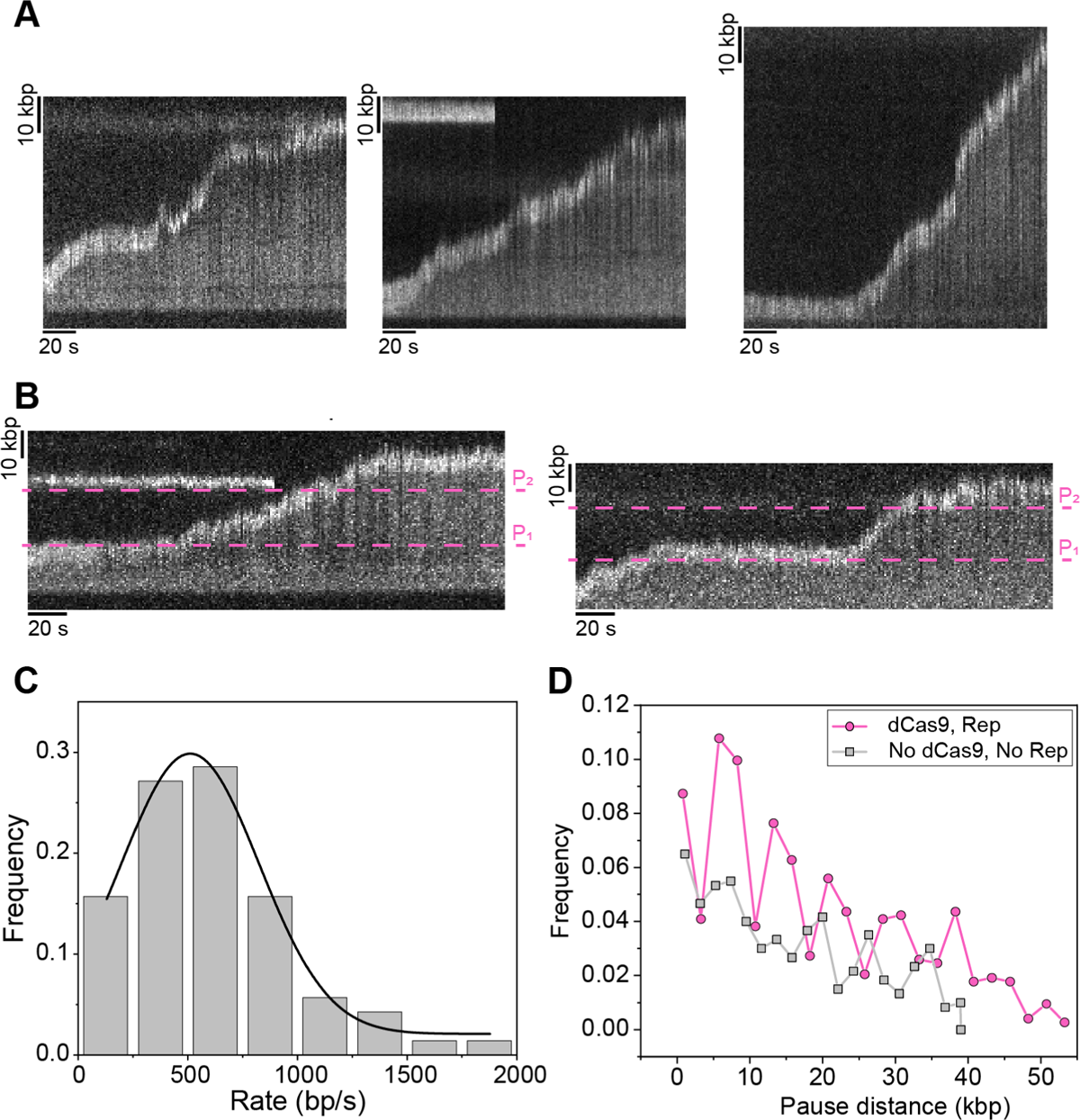
Single-molecule rolling-circle DNA replication of an 18-kbp DNA template. **(A)** Three example kymographs of elongating DNA replication of Sytox orange-stained 18-kbp rolling-circle DNA template (gray) (*n* = 32 molecules; replication efficiency of 6.4 ± 0.6% (S.E.M.)). Large circle of template is resolved at the tip of the replicating molecule at a higher intensity as it is stretched out by flow. **(B)** Two example kymographs of elongating DNA replication showing multiple pausing and rescue events by dCas9-cgRNA1 (0.25 nM) and Rep WT (10 nM) in solution (*n* = 26 molecules; 3.1 ± 0.2%). The dashed lines (magenta) indicate the theoretical pause start sites at approximately 17 kbp (P_1_) and 36 kbp (P_2_). **(C)** Histograms of the rate of replication for 18-kbp rolling-circle DNA templates (517 ± 129 bp s^-1^, *n* = 70) as in (A), fit to a Gaussian distribution. **(D)** Pairwise distance analysis of the paused start sites of 18 kbp replication rescue events in the presence of 10 nM Rep-AF647 and 0.25 nM dCas9-cgRNA1 (*n* = 37 pauses/733 kbp) (magenta) and absence (*n* = 28 pauses/600 kbp) (gray) for the first 60 kbp of DNA products.

**Supplementary Figure S9:**
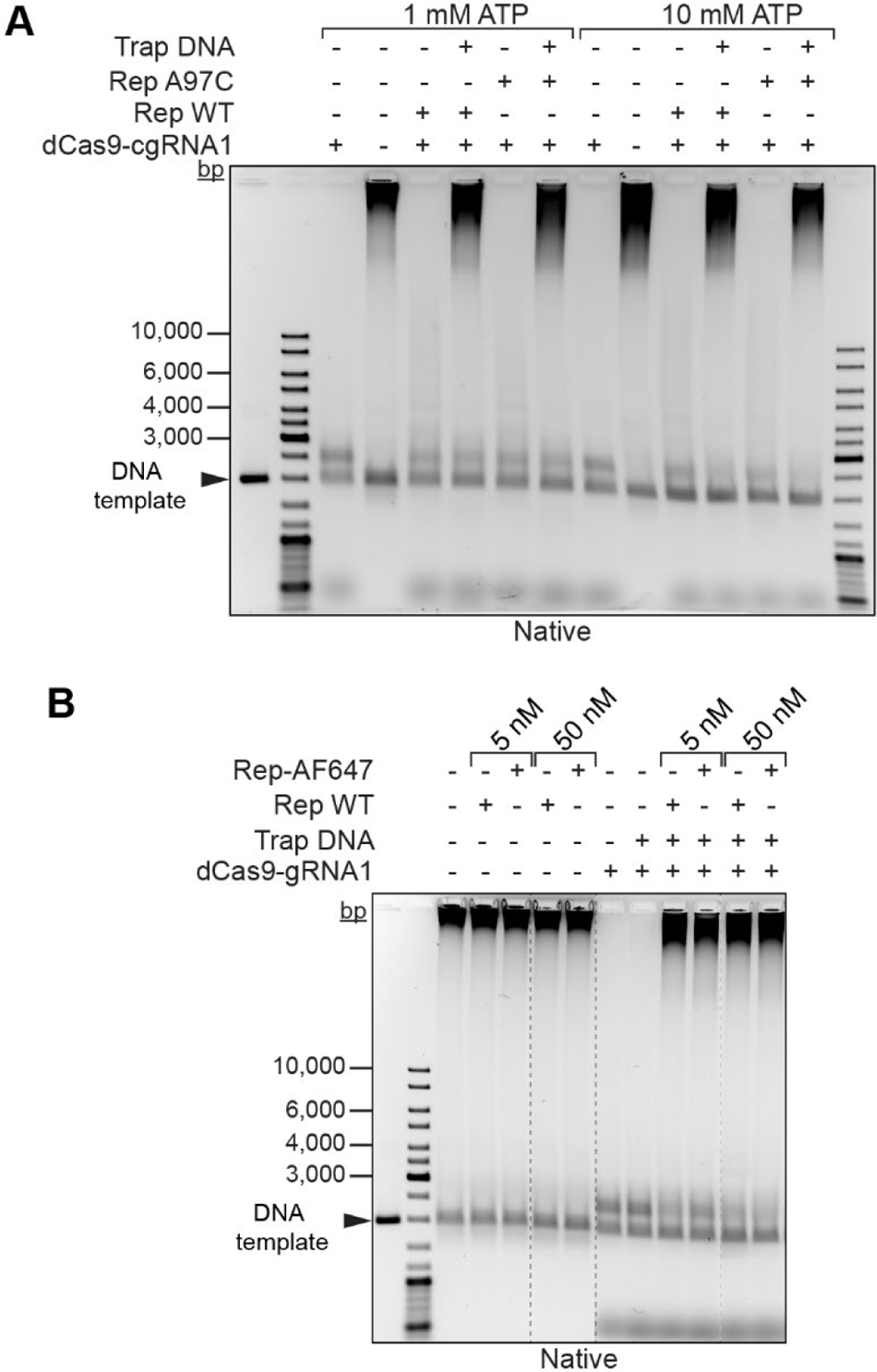
Ensemble replication rescue activity tests of Rep A97C and Rep-AF647. **(A)** Rep WT and Rep A97C (50 nM) efficiently remove dCas9-cgRNA1 (50 nM) complex from DNA templates and rescue stalled DNA replication. A 10-fold increase in ATP concentration (and 2-fold increase in MgCl_2_ concentration), results in a greater extent of DNA replication rescue, evident by decreased intensity of DNA band at approximately 2.5 kbp. **(B)** Rep-AF647 successfully removes dCas9-cgRNA1 complex from DNA templates. Irrelevant lanes to the figure are cropped out (dashed lines).

**Supplementary Figure S10:**
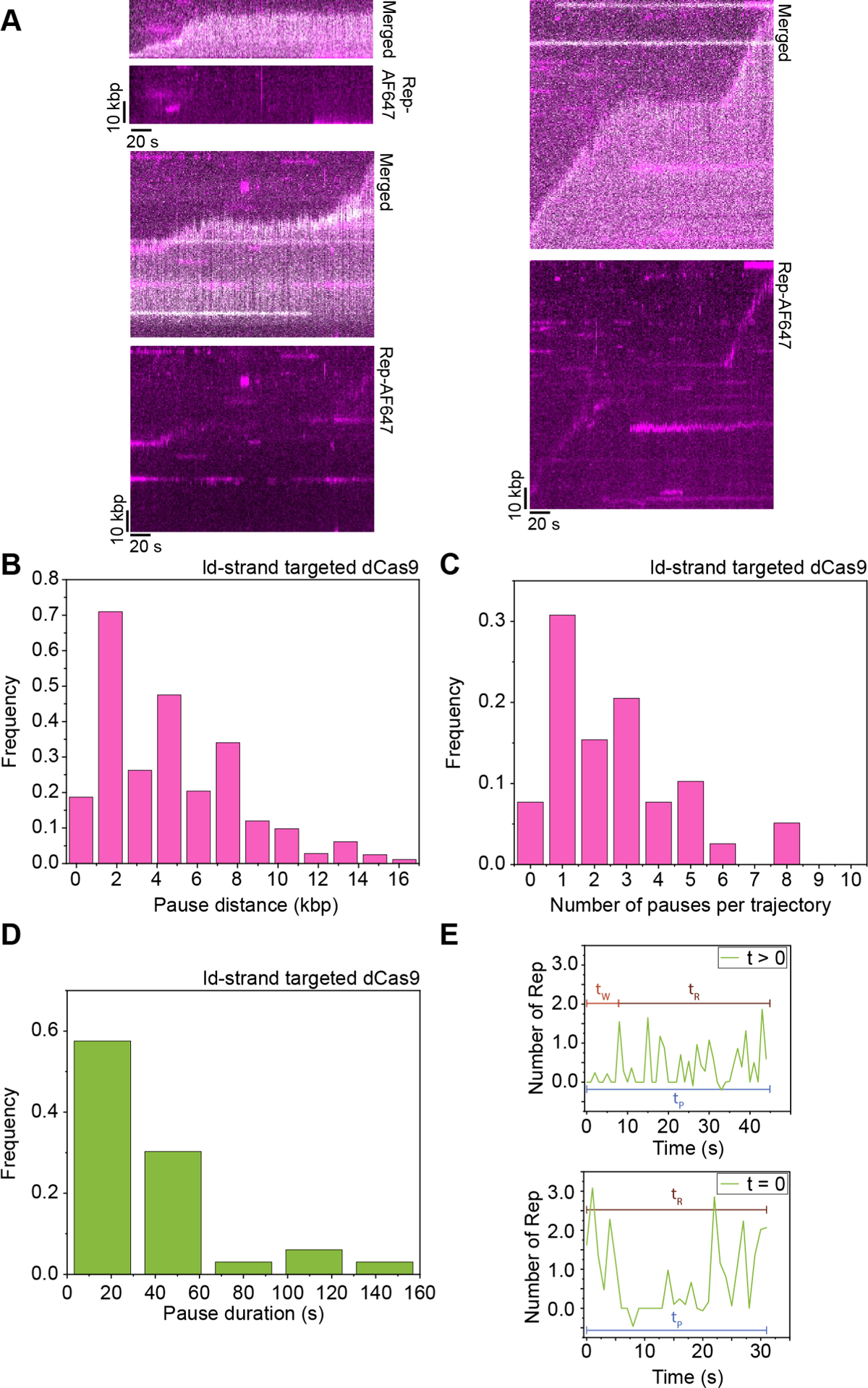
Single-molecule replication rescue of 2-kbp rolling-circle DNA templates. **(A)** Three example kymographs of rolling-circle DNA replication pausing and rescue events in reactions containing dCas9-cgRNA1 (0.25 nM) and Rep-AF647 (10 nM). Merged kymographs of Sytox orange-stained DNA products (gray) and Rep-AF647 (magenta) (top), and Rep-AF647 intensity alone (bottom). (replication efficiency of 2.9 ± 0.3%). (B–D) assays containing leading-strand target dCas9-cgRNA4 complexes (0.25 nM) and Rep-AF647 (10 nM) show **(B)** periodicity of pausing and rescue events (*n =* 43 pauses/358 kbp), **(C)** number of pausing events per replicating molecule (*n* = 40 molecules; replication efficiency of 2.0 ± 0.3%), and **(D)** mean duration of pauses of 38 ± 17 s (*n =* 33 pauses). **(E)** Annotated example of Rep-AF647 intensity traces over time showing the time points used for determining the pause duration (*t*_P_ – blue), association wait time (*t*_R_ – red) and pause resolve time (*t*_R_ – dark brown) for each Rep activity (*t* > 0, left; *t* = 0, right).

**Supplementary Figure S11:**
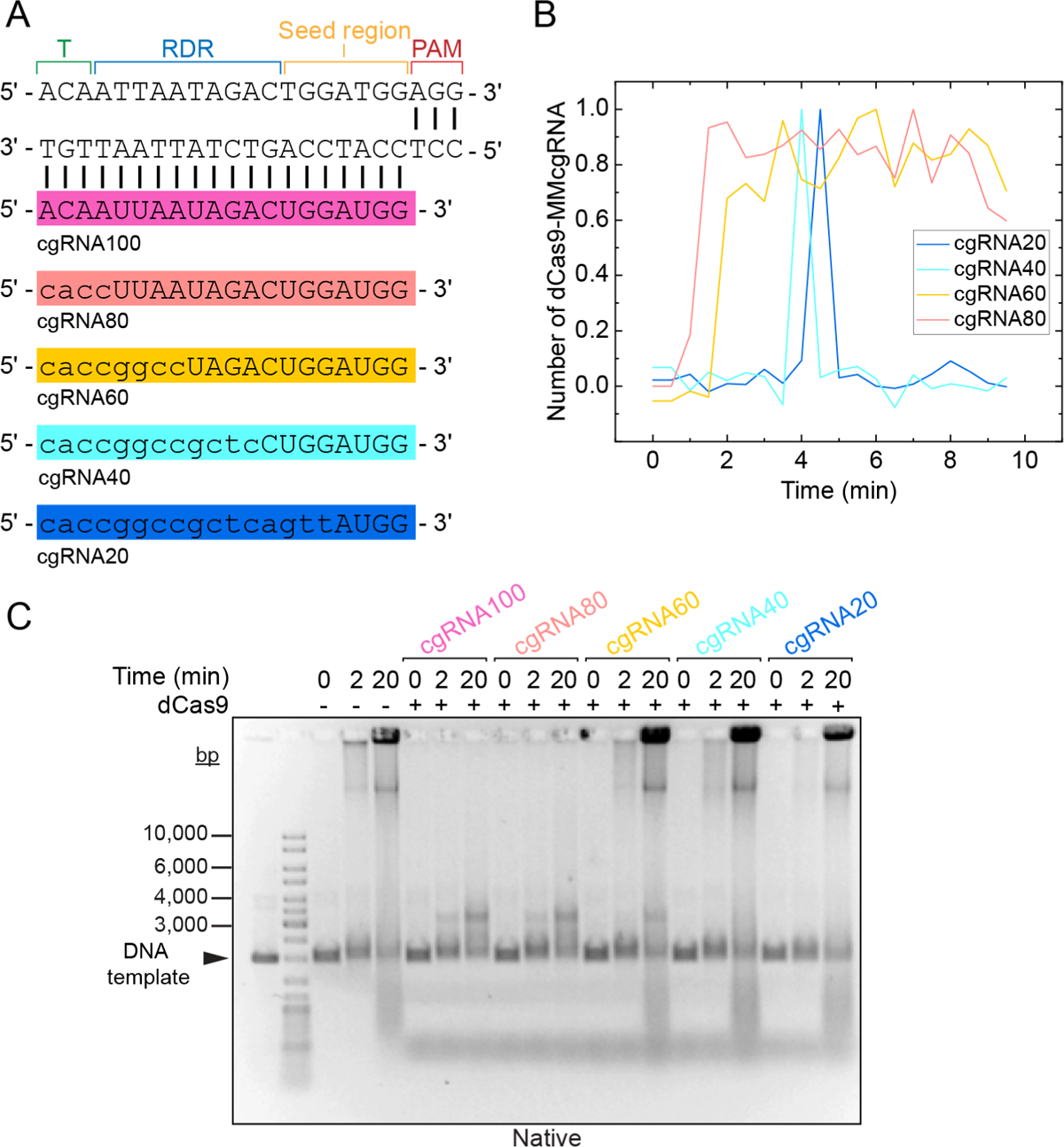
Characterization of dCas9 complexed with mismatch cgRNAs. **(A)** Designs of cgRNA1 and various mismatch gRNAs ranging from 80–20% complementarity to the target sequence of cgRNA1 (or cgRNA100). Mismatches, denoted by lowercase letters, span across the DNA-RNA hybrid from the PAM distal region (or terminal region (T – green)), reversibility-determining region (RDR – blue), seed region (yellow) and to the PAM proximal region (red). **(B)** Example lifetime intensity trajectories of dCas9-cgRNA80 (peach), dCas9-cgRNA60 (yellow), dCas9-cgRNA40 (light blue) and dCas9-cgRNA20 (blue). Intensity is corrected for the intensity of a single dCas9-cgRNA-Atto647 molecule measured by photobleaching analysis. **(C)** Time course ensemble characterization of dCas9-cgRNAMM complexes blocking rolling-circle DNA replication over 20-minutes.

